# Statistical modelling of aquatic size spectra: integrating data from multiple taxa and sampling methods

**DOI:** 10.1101/2022.08.29.505693

**Authors:** Henrique Corrêa Giacomini, Derrick T. de Kerckhove, Victoria Kopf, Cindy Chu

## Abstract

Size spectra are used to assess the status and functioning of marine and freshwater ecosystems worldwide. Its use is underpinned by theory linking the dynamics of trophic interactions to a power-law decline of abundance with body size in ecological communities. Recent papers on empirical size spectrum estimation have argued for Maximum Likelihood Estimation (MLE) of power-law probability distributions as a more accurate alternative to traditional linear regression approaches. One major limitation of currently used size spectrum estimators from MLE is that they cannot account for the use of multiple sampling protocols, nor the distortions caused by gear size selectivity, and therefore they become restricted to a relatively narrow taxonomic group and size range. Further progress in the field requires new methods that are flexible enough to combine multiple trophic groups and sampling gears into a single size spectrum estimate, while taking advantage of more accurate distributional approaches. The method we propose in this paper fills this gap by deriving the distribution of observed sizes explicitly from the underlying power-law spectrum and gear selectivity functions. It specifies likelihoods as a product of two components: (i) the probability of belonging to a given group and (ii) the probability distribution within the group. Using Bayesian estimation, we applied the method to surveys of phytoplankton, zooplankton, and fishes in lakes of Quetico Provincial Park, northwestern Ontario, using Van Dorn samplers, zooplankton nets, gillnets, and hydroacoustics. The results show that the spectra estimated from subsets of trophic groups or gears are weak predictors of more complete spectra, highlighting the importance of using more inclusive community data. The two-component partitioning of likelihoods also helped demonstrating the existence of *between-group* spectrum slopes that were overall steeper than *within-group* slopes, indicating that heterogeneity of trophic transfers across the size spectrum is an important factor structuring these ecosystems.

## 1. Introduction

In aquatic communities, predators are larger and less abundant than their prey (Elton, 1927; Cohen et al., 2003). This negative relationship between body size (*y*, representing weight or length) and abundance (*n*, in number of organisms per habitat area or volume, per unit weight or length) can be summarized as a size spectrum: *n* = *κy*^−*λ*^, a power law predicted by theory to result from predation dynamics in size-structured communities (Kerr and Dickie, 2001; Andersen et al., 2016). When represented on a logarithmic scale, the spectrum becomes a straight line with intercept given by log (*κ*) and slope given by −*λ* (a reason why this parameter is traditionally referred to as *size spectrum slope*, despite being an exponent). The intercept is an indicator of total abundance or productivity (Boudreau and Dickie, 1992; Sprules and Barth, 2016) while the slope represents the rate of abundance decline with size, which is linked to energy transfer efficiency, predator-prey size ratios, metabolism, and mortality, amongst other size-dependent processes (Andersen and Beyer, 2006; Trebilco et al., 2013; Mehner et al., 2018). Given its simplicity and theoretical support, size spectrum slopes have been used by many studies to quantify the impacts of perturbations such as pollution, warming, nutrient change, habitat modification or fisheries exploitation in marine and freshwater communities (e.g., Sprules and Munawar, 1986; Rice and Gislason, 1996; Emmrich et al., 2011; Yvon-Durocher et al., 2011; Atkinson et al., 2021).

To be a consistent indicator of ecosystem process and responses to perturbations, the size spectrum slope must cover a large enough range of sizes in the sampled community, which may involve the inclusion of multiple taxonomic or trophic groups, such as phytoplankton, zooplankton, macroinvertebrates, and fish (Sprules and Barth, 2016). When sampling diverse taxonomic groups from the same community, surveys are most likely to rely on the use of different gear (e.g., net hauls versus hydroacoustics) or configurations (threshold detection frequencies in hydroacoustics), as each gear is selective with respect to habitat, taxon, and body size. The traditional approach for size spectrum inference from community data is to apply a linear regression between log-transformed and binned body size values against log-transformed abundance or biomass (Sprules and Barth, 2016). This method is flexible enough to accommodate multiple taxonomic groups from different sampling gears in the same analysis, as it involves a simple transformation of the response variable (e.g., dividing catch by sampling efforts and known gear selectivity functions). More recently, the linear regression approach has been criticized for not recognizing that body size samples are in fact frequency distributions (White et al., 2008; Edwards et al., 2017). By representing data as a bivariate set of a few size-abundance coordinates, it does not properly weight individual size observations as the real sampling units, which leads to poor inference. The recommended alternative is to adopt a more explicit distributional approach, one that estimates the exponent of the underlying power-law distribution from the likelihoods of individual observations. Within this approach, the method of choice by previous papers has been the Maximum Likelihood Estimaton (MLE) of the Pareto type 1 or truncated power-law distributions. It gives more accurate and precise estimates than regression-based methods (White et al., 2008; Edwards et al., 2017, 2020) and has already been used by recent empirical studies (e.g., Robinson et al., 2017; Braun et al., 2021). The currently used size spectrum estimators from MLE are nevertheless quite limited. The main reason is that they assume the data are free from distortions caused by gear size selectivity or by variable sampling effort, and thus can only be applied to taxonomic groups sampled by a common survey protocol and within relatively narrow size limits. Another problem is that they are not easily integrated into more complex statistical model structures, which still prevents them from being fully adopted (e.g., Heather et al., 2020; Perkins et al., 2021; but see Robinson et al., 2017).

The method we describe in this paper models body size data explicitly as frequency distributions resulting from selective gear sampling of an underlying community size spectrum. It allows for the integration of data from multiple taxonomic groups and sampling protocols, thus overcoming the limitations of current statistical approaches. Through Bayesian estimation, we applied the method to community surveys of six inland lakes in Quetico Provincial Park, northwestern Ontario, Canada. These surveys include three trophic groups (phytoplankton, zooplankton, and fish) from the pelagic zone, in addition to fish sampling by benthic gillnets using two distinct protocols. Given that doing comprehensive community surveys can incur high logistic and sample processing costs, there was also an interest in verifying whether the spectra from subsets of the community can still serve as reliable indicators of fuller community spectra. So, a second objective was to test whether slopes estimated from different trophic groups and survey methods provide consistent indicators of size spectrum variation among lakes.

## 2. Methods

### 2.1. Sampling and data processing

A detailed description of sampling methods and data processing is given in the Supplementary Material A. Six lakes ranging from 233 to 2484 ha in size with maximum depths of 26.6 to 71 m were sampled between July and August 2016 and 2017 for this study. The lakes are located within Quetico Provincial Park, northwestern Ontario, Canada (centroid of park ~ 48.4°N, −91.4°W). Size spectra have been adopted in the Fisheries Management and Aquatic Ecosystem Stewardship Plan for the park as one of the indicators used to evaluate the biological status of lake communities (Ontario Parks, 2019). Phytoplankton, zooplankton, and fish were sampled following the Broad-scale Monitoring (BsM) program of inland lakes in Ontario (Lester et al., 2021), using Van Dorn samplers, zooplankton nets, and gillnets, respectively. Additional hydroacoustic surveys for fish were carried out within 1 to 2 weeks of gillnetting on each lake. Because phytoplankton, zooplankton, and hydroacoustic fish samples allowed for absolute estimates of organism densities (number of individuals per m^3^), they could be combined to estimate a common size spectrum and are thereafter referred to as *community* data. The *gillnet* fish data consisted of samples from two gillnetting standards, as per BsM protocol: the North American (NA), with relatively large mesh sizes, and the Ontario (ON) standard, with smaller mesh sizes. Due to the lack of known catchability for both gears and of marked fish populations allowing to estimate absolute population sizes, the gillnet data could only be combined among themselves but not with the community data, so they were used to provide a separate set of size spectrum estimates for fish. A broader dataset from the BsM, consisting of mesh-specific catch data from 99 lakes, was used in addition to estimate and compare gillnet selectivity models for fish (Supplementary Material B).

### 2.2. Analysis

The most important assumption is that the size distribution of organisms in a community can be described by a power function:

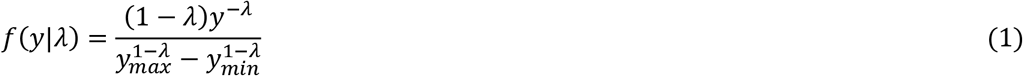

where *y* is a size variable (e.g., length or weight) defined within given limits (*y_min_*≤*y* ≤ *y_max_*), and *f*(*y*|*λ*) is the probability density function of *y*, conditional on its parameter *λ*, which is the magnitude of the size spectrum slope (the slope itself is −*λ*), with the condition that *λ* ≠ 1. Our general approach for estimating *λ* is to derive a probability density function of body size that is the product of two components: (i) the probability density function of size within a given group *g* (trophic group or gillnet gear), denoted here by *o_g_*(*y*|*λ, **θ**_g_*), where ***θ**_g_* is the set of parameters determining the gear- or group-specific size selectivity function; and (ii) the probability that an observed individual belongs to group *g* (i.e., the probability mass function of the discrete variable *g*), denoted by *ρ*(*g*|*λ, **θ, v***), where ***θ*** = {***θ**_g_; g* = 1,2,..} is the set of all selectivity parameters and ***v*** = {*v_g_*; *g* = 1, 2,..} is the set of sampling efforts associated with the groups or gears, measured as e.g., water volume (m^3^) in hydroacoustic surveys or total length of nets (m) in gillnet surveys. They both combine the underlying size spectrum *f*(*y*|*λ*) with a selectivity function *s_g_*(*y, **θ**_g_*), which measures the probability of detecting or catching any individual of size *y* per unit sampling effort. The general expressions for these functions are:

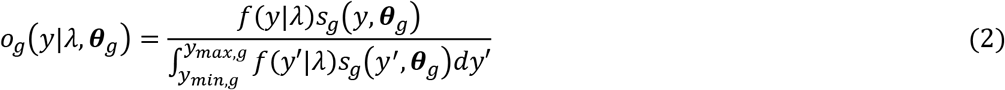

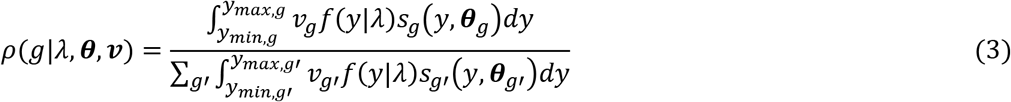

where *y_min,g_* and *y_max,g_* are the group- or gear-specific minimum and maximum sizes. The derivation of these and more specific probability functions used for the community and gillnet data analyses is explained in the Supplementary Material C. The likelihood associated with *λ* for each individual observation *i* is given by the product of Equations (2) and (3), i.e., *L*(*λ*|***θ**_g_, y_i_, g_i_, **v***) = *ρ*(*g_i_*|*λ, **θ, v***)*o_g_*(*y_i_*|*λ*, ***θ**_g_*). With the likelihood function and a prior distribution for *λ*, Bayesian methods can then be applied to estimate the posterior distribution conditional on the observed data and known selectivity parameters. If the selectivity parameters are not known but are jointly estimated with *λ*, then another likelihood function is necessary, i.e., *L*(***θ**_g_*|***X***). in this case associated with ***θ**_g_* and additional gear-specific data ***X*** (such as a matrix with individual body size and mesh information from gillnet catches). With both likelihood functions and priors for *λ* and ***θ**_g_*, a joint posterior *p*(*λ, **θ**_g_*|***y, g, X***) can be estimated considering uncertainty in both the size spectrum and the gear selectivity parameters.

In usual size-spectrum surveys, each individual observation *i* can be assigned to both a group category *g_i_* and a body size *y_i_*, so the total log-likelihood is:

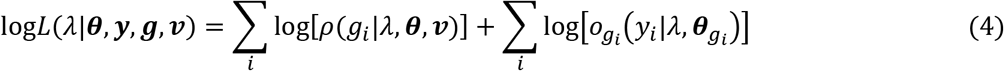

which is conveniently partitioned between the two probability components. This can be useful for two reasons. Firstly, if the observed data does not contain information on either individual group identity, sampling effort or even individual body size values, the size spectrum slope *λ* can still be estimated (provided the selectivity parameters are known or estimated with an additional dataset ***X***). If, on the one hand, we do not have individual size information, but only the total number of observations and sampling effort for each group, the log-likelihood becomes log*L*(*λ*|***θ, g, v***) and simplifies to the first summation term of Equation (4). If, on the other hand, we ignore information on group-specific sampling efforts, the log-likelihood becomes log*L*(*λ*|***θ, y, g***) and simplifies to the second summation term. Still another possibility is if we know individual sizes and sampling efforts but have no information on the group each individual belongs to. The log-likelihood in this case, denoted by log*L*(*λ*|***θ, y, v***) will be based on a mixture distribution given by the sum of *ρ*(*g*|*λ, **θ, v***)*o_g_*(*y*|*λ, **θ**_g_*) across groups (Supplementary Material C)

The second reason this partitioning is useful is that differences between *λ* estimated from different likelihood components can provide insights into how the size spectrum is structured in terms of its within-versus between-group variation. In principle, all likelihood functions described above should give similar estimates of *λ*, provided the community size spectrum strictly follows Equation (1). However, this will not be true in many situations in which the size spectrum has a secondary structure in addition to the primary, power-law structure. For instance, if trophic transfers occur more frequently between groups than within a group (e.g., fish eats zooplankton more often than other fish), a greater fraction of energy is lost from one group to another due to the inevitable inefficiency of energy conversion from prey to predator (i.e., there is a greater deviation from the “energy equivalence” expected within single trophic level, Trebilco et al., 2013). The expected result is that the *between-group λ* estimated through *ρ*(*g*|*λ, **θ, v***) will be higher (more negative slope) than the *within-group λ* estimated through *o_g_*(*y*|*λ, **θ**_g_*).

The general approach outlined above was applied separately to the community and gillnet data. The most distinctive aspect of analyzing gillnet data is its more complex selectivity function, which depends on the combination of mesh sizes. Our selectivity analysis introduces two novel aspects: (i) it proposes a new function for retention selectivity called *Symmetric Exponential* and (ii) it uses body weight to derive a modified size metric that better accounts for species differences in body shape. Both aspects were shown to greatly improve model fit to the data (see Supplementary Material B). After specifying the form of *s_g_*(*y, **θ**_g_*), we derived analytical solutions for the observed size distributions *o_g_*(*y*|*λ, **θ**_g_*) and group-specific probabilities *ρ*(*g*|*λ, **θ, v***), which in turn provided the basic ingredients for the likelihood functions and the estimation of *λ*. Bayesian estimation was carried out in MATLAB R2020b using the slice sampling MCMC algorithm (Neal, 2003) (see Supplementary Material D for details on all Bayesian analyses).

We estimated the slope of several size spectrum components, following two criteria: (i) based on different combinations of groups or (ii) based on different components of the likelihood function (Equation 4). For the community data, the first criterion used three combinations of trophic groups, from more to less inclusive: phytoplankton + zooplankton + fish (PZF slope, available for three lakes only), zooplankton + fish (ZF slope), and fish only (F slope). For the gillnet data, the groups were based on gear types: NA slope, ON slope, and BsM slope (combining both gears). In all these cases, the slope estimation was based on the complete likelihood function. In the second criterion, the slope was estimated using the first component of the likelihood function (Between-group slope) or the second component (Within-group slope), in both cases including together all groups (trophic groups or gillnet gears) available in each lake.

## 3. Results and Discussion

The slopes showed little consistency across different trophic and gear group combinations (Figure 1). One exception is the ZF versus ON slope (Figure 1H), whose correlation (r = 0.97, p = 0.001) is significant at 5% even after a sequential Bonferroni correction for multiple tests. A possible reason is that ON gillnets catch relatively small fish, which are near in size to the spectrum connecting fish to zooplankton. However, this does not explain why other supposedly close spectra, such as ZF and F, or any combination involving fish only (e.g., BsM vs F), did not have correlated slopes. Even though the data are quite limited, including only six lakes (and only three with phytoplankton information), the slopes should have been all strongly correlated if the spectra strictly and consistently followed a power-law, regardless of group or habitat (i.e., benthic versus pelagic).

**Figure 1.**
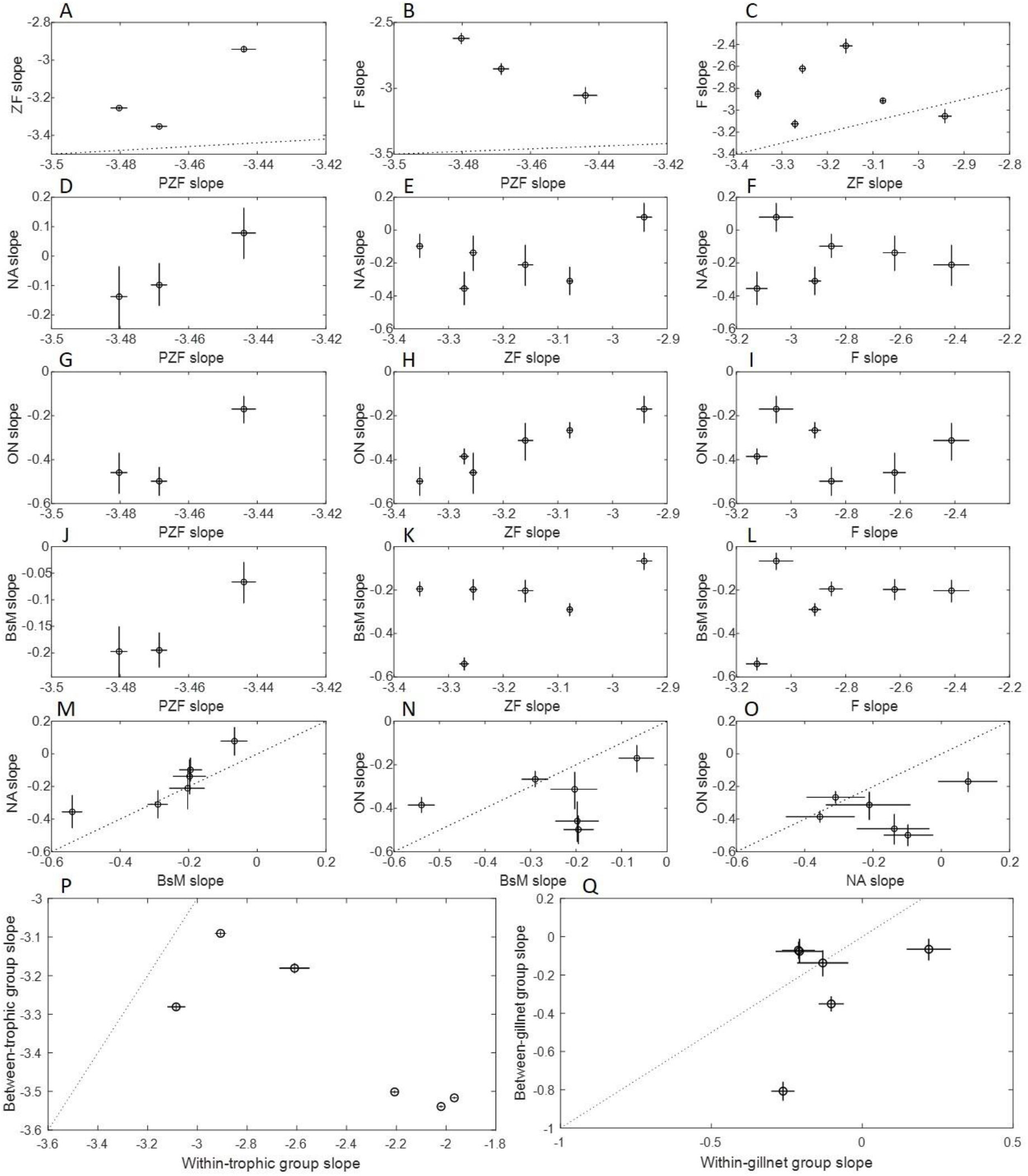
Size spectrum slopes (median and 95% credible intervals) estimated for different combinations of trophic groups and gillnet gears (A-O), or likelihood components (P-Q). The initials P, Z, and F represent phytoplankton, zooplankton, and fish from the community data, grouped from less inclusive (F only) to more inclusive (PZF). The gillnet data are represented by three gear combinations: North American standard (NA) only, Ontario standard (ON) only, or both together making up the Broad-scale Monitoring protocol (BsM). The dotted lines are 1:1 lines. P-Q plot the slopes estimated from the *between-group* likelihood component (Equation 4) against slopes estimated from the *within-group* component. P: groups were defined by trophic groups from the community data; the three points at the bottom-right end are the lakes with data for all groups. B: groups were defined by gear types from the gillnet fish data.

The overall lack of association among groups indicates that the size distributions in these communities are more complex, with a pronounced secondary structure. This is evident from the observed size distributions, which deviate systematically and non-linearly from the power-law expectation (Figures E1–E4 in the Supplementary Material E). Such deviations can arise from several processes hypothesized to affect the secondary structure, including: (i) trophic cascades (Rossberg et al., 2019), (ii) non-equilibrium dynamics caused by inherent instability (Law et al., 2009), fishing (Rochet and Benoit, 2012), or seasonal pulses of primary production and spawning (Datta and Blanchard, 2016), (iii) uneven distribution of life history strategies across the size range (Andersen et al., 2016), (iv) uneven distribution of energy subsidies (Perkins et al., 2021), (v) uneven distribution of trophic interactions across the size range (Chang et al., 2014) or habitats (Blanchard et al., 2009). This multitude of processes makes it difficult to pinpoint the actual causes of observed deviations in this limited sample of lakes. However, it is likely that uneven trophic interactions play a major role, for two reasons. Firstly, the slopes from the community data get more negative for more inclusive groupings, i.e., PZF < ZF <F. This implies that adding trophic groups leads to disproportional jumps in abundance, which in turns steepens the mean spectrum slope. Secondly, the slopes based on between trophic group variation were steeper than those based on the within-group variation (Figure 1P-Q). The difference was even more pronounced for lakes that included all three trophic groups (Figure 1P). For the gillnet data, the between- and within-gear slopes were more similar (Figure 1Q), which is expected as they represent a single trophic group (fish). These results are consistent with the hypothesis that predators from one group, such as fish, feed predominantly on organisms from another group, such as zooplankton: with relatively fewer trophic transfers within a group than between, less energy is lost due to transfer inefficiencies and more biomass can be retained at larger sizes, therefore resulting in shallower within-group slopes. The results are in line with equilibrium size spectrum theory (Trebilco et al., 2013) and with the formation of group-specific “domes” in size distributions (Boudreau et al., 1991; Yurista et al., 2014). They highlight that including a wide range of trophic groups is essential if the size spectrum is to be used as a consistent indicator of energy transfers in ecosystems.

Another noticeable pattern is that the fish slopes estimated from the hydroacoustic data were more negative than the estimated from the gillnet data (F slopes varied from around −3.2 to −2.4 whereas NA, ON, and BsM slopes varied from around −0.6 to 0.1). This could be partially due to ecological differences between habitats these gears survey (pelagic versus benthic) but are more plausibly explained by methodological factors. The hydroacoustic slopes were based on body length whereas the gillnet slopes were based on weight, so the first are expected to be in average 3 times higher in magnitude (more negative) than the second if a general cubic weight-length relationship applies. Another reason is that the gillnet selectivity model can only account for mesh-specific variation in retention rates, conditional on fish encountering the net (Anderson, 1998). Ignoring encounter rates can bias slope estimates because fish movement – which determines the chance of encountering the net – is also dependent on body size. Movement rates in general increase as a power function of size, scaling as *y^β^* (Rudstam et al., 1984), so the size distribution of fish encountering the nets should be proportional to *y^−λ+β^* instead of the assumed *y^−λ^*. For this reason, the gillnet slopes should be interpreted as “apparent slopes”, shifted up from the true value by a fixed amount *β*. For instance, if movement rate scales with body weight to the power *β* = 0.13 (Andersen and Beyer, 2006), a slope estimated as 0 from gillnet data would actually represent a true spectrum slope of −*λ* = −0.13.

The gillnet selectivity analysis led to two important developments. The first comes from the observation that catch rates decline in an approximately symmetrical and exponential way as the body size/mesh size log-ratio deviates from its optimal value. This resulted in the Symmetric Exponential model, which provided the best fit to our data and can serve as a useful alternative to commonly used selectivity models (Millar and Fryer, 1999). The second is the recognition that fish from different species can vary markedly in body shape, thus raw length values can be poor predictors of catch vulnerability by gillnets. Here we used a modified size variable called *Effective Length (EL*), which is derived from body weight (*W*) by imposing an isometric transformation *EL* = *W*^1/3^. For any given raw body length, fish species with more elongated body, such as Lake Trout (*Salvelinus namaycush*), are expected to be vulnerable to relatively smaller mesh sizes when compared to species with “deeper” body, such as Smallmouth Bass (*Micropterus dolomieu*), a difference that is well captured by their smaller *EL*. This might not be so relevant in usual single species applications but can make a difference within the multispecies context of empirical size spectrum studies, which to date have been based on the use of minimum size thresholds as a surrogate for gillnet selectivity (e.g., Emmrich et al., 2011).

In conclusion, the analytical method proposed in this paper – and applied to inland lake communities of Quetico Provincial Park – provides a way of integrating body size data from multiple taxonomic or functional groups and sampling protocols. It combines the flexibility of linear regression approaches with the statistical rigor of distributional approaches. There are two important ways in which the method can be expanded to tackle more general questions in size spectrum research. One is incorporating environmental and anthropogenic covariates to test for hypothesis about size spectrum variation. This can be done within a Bayesian framework by modelling each size spectrum parameter (e.g., *λ* or the total community abundance) explicitly as a function of these covariates and adding them to the network of conditional probabilities determining the expected distributions of observed data (i.e., *o_g_*(*y*|*λ, **θ**_g_*) and *ρ*(*g*|*λ, **θ, v***)). Another important direction for further research is to develop probability functions that describe the secondary structure of size spectra. A promising possibility is to use a wave function to account for regularly spaced fluctuations around the primary power-law trend, similar to the empirical model used by Rossberg et al. (2019) but applied more generally. Regular fluctuations in the size spectrum are predicted by theory to occur from several processes, not restricted to Rossberg et al. (2019) top-down cascade mechanism, but including non-equilibrium dynamics (Law et al., 2009), fisheries-induced cascades (Rochet and Benoit, 2012), and seasonal pulses (Datta and Blanchard, 2016). A glimpse of these fluctuations can be seen in the size distributions of our study lakes (Figure E1–E4), although the limited amount of data prevented estimation of a more complex model. Characterizing fluctuations by a well-defined function and interpretable parameters will be useful to disentangle the effect of different factors, such as predator/prey size ratio affecting their period or primary productivity affecting their amplitude (Rossberg et al., 2019). The last few decades have seen the development of theoretical models explaining the variation in structure and dynamics of aquatic size spectra, as well as of survey techniques covering a broad range of taxonomic groups and habitats. Further progress in the field will also require the development of statistical methods capable of connecting existing theory to data.

## Acknowledgements

The authors are thankful to the Broad-scale Monitoring science team and all crew members who collected data and made this work possible. Abby Daigle and Lee Gutowsky helped in the field with hydroacoustic surveys. Funding for this research was provided by the Ministry of Natural Resources and Forestry’s Special Purpose Account.

## Supplementary Material

### Supplementary Material A: sampling methods and data processing

#### A1. Study area and field collection

Quetico Provincial Park in Northwestern Ontario, Canada (centroid of park ~ 48.4°N, −91.4°W), contains 4718 km^2^ of remote wilderness area which comprises over 2000 lakes in near pristine condition, of which a subset is monitored under the provincial Broad-scale Monitoring Program (BsM, Lester et al., 2021). Six lakes ranging from 233 to 2484 ha in size with maximum depths of 26.6 to 71 m were sampled for this study: Camel, Louisa, Mack, McEwen, Saganagons, and Wolseley.

Phytoplankton, zooplankton, and fish community data were collected between July and August 2016 and 2017 in the six study lakes to meet the objectives of this study. Wolseley, McEwen and Saganagons lakes were surveyed in July-August 2016 while Camel, Mack and Louisa lakes were surveyed in July-August 2017. Phytoplankton and zooplankton were sampled as part of the Broad-scale Monitoring (BsM) protocol for Inland Lakes (Sandstrom et al., 2013). For the phytoplankton collection, Secchi depths were measured and the entire euphotic zone (twice the Secchi depth) was sampled using a combined integrated water sample (Schneider et al., 1983). To produce the integrated sample, Van Dorn samples (2 L of water) were collected 1 m below the surface and at 2 m depth intervals until the euphotic zone was sampled. A 250 ml sub-sample was collected for each depth and combined in a 4 L jug. A 500 ml sub-sample was collected from the jug and preserved with Lugol’s for laboratory analysis (see below for more details). Zooplankton samples were taken using vertical hauls from the deepest location in each lake as described in the manual’s Appendix XV (Sandstrom et al., 2013). The 63 μm zooplankton nets were hauled a rate of 1 m per second through the entire column. Net hauls were repeated three times with at least 15 min between hauls to allow the zooplankton to mix in the water column. Zooplankton collected in the cod end were preserved using Lugol’s in 500 ml or 1L jar for laboratory analysis.

Fishes were sampled using two methods: (i) the BsM gillnetting protocol and (ii) through hydroacoustic surveys, which were completed within 1 to 2 weeks of gillneting on each lake. Each method is described in more detail below. Because only hydroacoustic, zooplankton, and phytoplankton samples allowed estimation of absolute densities (organisms/m^3^), they could be grouped in the same analyses to estimate a common community size spectrum and are hereby referred to as *community data*. The gillnet samples could not provide estimates of absolute fish density (due to the absence of marked populations or a known catchability coefficient), so could not be combined with the rest of the community data, and for this reason were analyzed separately and are hereby referred to simply as *gillnet data*.

##### A1.1. Broad-scale Monitoring gillnet surveys and data processing

The Broad-scale Monitoring (BsM) protocol is described in detail by Sandstrom et al. (2013). Two gillnet standards, the North American (NA, with relatively large mesh sizes) and the Ontario (ON, with relatively small mesh sizes) were deployed in a random depth-stratified manner. Both gears were fished in the following depth strata: 1–3, 3–6, 6–12, 12–20, 20–35, 35–50, 50–75 m. The number of NA net gangs ranged from 4 to 555 (24 to 160 in our six study lakes) per survey, depending on lake size; the number of ON net gangs ranged from 0 to 140 (22 to 64 in our study lakes). ON net gangs were 12.5 m in length, 1.8 m high, with five panels of mesh ranging from 13 to 38 mm in stretched mesh (i.e., 13, 19, 25, 32, and 38 mm). NA net gangs are 24.8 m in length, 1.8 m high, with eight panels of mesh ranging from 38 to 127 mm (38, 51, 64, 76, 89, 102, 114, and 127 mm). Nets were set in the afternoon and lifted in the morning with a target soak time of 18 h. A double gang strap was used (i.e., two nets strung together). Fish were identified to species and their lengths and weights measured and recorded.

The total BsM data included surveys from 2007 to 2017 in 859 lakes. For the purpose of our analyses, we used two subsets: (i) data with recorded mesh size information (*mesh-specific data*), and (ii) data from the six study lakes in Quetico Park (*Quetico data*). The mesh-specific data included fork-length (mm), weight (g), and mesh size (mm) from individual fish catches (total 32755 individuals from 99 lakes). The Quetico data included fork-length and weight of 3986 fishes from the six lakes. We used the mesh specific data to estimate and compare gillnet size selectivity models (Supplementary Material B), whereas the Quetico data were the focus of this study for size spectrum estimation.

As per BsM protocol, when the catch from an ON net included numerous small fish and prevented timely processing of every individual, body sizes were assigned to 10mm wide fork length classes (10-20mm, 20-30mm, …). They included 13460 fishes (41%) from mesh-specific data and 2448 (61%) from the Quetico data. For these fish, we individually assigned body weight values based on the mid-point of their fork length class and on species-specific length-weight relationships estimated directly from the BsM data or from known relationships for the species in Ontario.

##### A1.2. Hydroacoustic Surveys and data processing

Mobile hydroacoustic surveys were completed using a mid-ship mounted pole and plate submerged at 0.3m with a Simrad Combi 13° x 21° 38 kHz/ 7° 200 kHz split-beam transducer (Kongsberg Maritime, Kongsberg, Norway) oriented vertically downward with a Simrad EK80 WBT and processing computer onboard. Simrad EK80 Echo sounder software was used to run the system at maximum ping intervals and parameter settings (Kongsberg Maritime, Kongsberg, Norway; Table 1). Transducers were calibrated in the field prior to surveys.

All acoustic data from the 38 kHz transducer were analyzed using Echoview® (Myriax Software Pty. Ltd. version 11.0.255.39592 64-bit edition) according to recommendations in the Echoview software manual and the standard operating procedures for the Great Lakes (Parker-Stetter et al., 2009); data analysis parameters are listed in Table 1. Erroneous data regions resulting from electronic noise and bottom detections were removed from the analysis area of echograms (Parker-Stetter et al., 2009). Bottom exclusion zones were set at 0.75 m above bottom to remove the acoustic dead zone and surface exclusion zones were set at 5.0 m below surface to remove the near-field zone and ring-down (Parker-Stetter et al., 2009). Analyses were restricted to areas with a minimum lake bottom depth of 10 meters to select for the pelagic zone only. Echoview operator functions were used for the following tasks, with either default parameters of those identified in Table 1: background noise removal, school detection, single target detection (note that method 2 was used) and fish tracks identification. We used an elementary distance sampling unit (EDSU) of 100 m horizontal distance by one-meter vertical depth bin to export and analyze acoustic data (de Kerckhove et al., 2015). Volume backscattering (Sv) and fish track target strength data (TS) were exported by EDSUs from Echoview. A minimum TS threshold of −46 dB was set to align with minimum body size requirements for data analyses. Echo integration was used for estimating abundance and biomass rather than echo counting to prevent target-coincidence biases (Soule et al., 1995; Parker-Stetter et al., 2009). Echoview exports were uploaded into the R v3.6.1 programming platform (R Core Team, 2018) for further data analyses.

Total lengths (mm) for fish were calculated from in situ TS data of individual fish targets using Love’s (1971) target strength model:

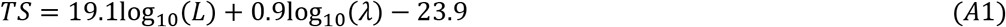

where *L* is fish length in meters and *λ* is the acoustic wavelength. TS data from adjacent one-meter layers were used to fill in missing layer data where no TS data were collected. TS data were grouped into ¼-log_2_ bins and used to parse S_v_ to determine target (fish) density by EDSU. Echo integration was completed to estimate fish density:

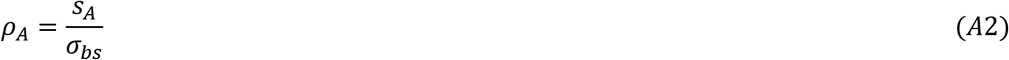

where *ρ_A_* is the estimated fish density (number/m^2^), *s_A_* is the average backscattering coefficient (m^2^/m^2^), and *σ_bs_* is the expected backscattering cross-section (m^2^).

**Table A1.** Simrad EK80 38 kHz hydroacoustic system settings and analysis parameters for 2016 and 2017 Quetico PP mobile surveys.

| Acoustic Survey Parameters |  |
| --- | --- |
| Frequency (kHz) | 38 |
| Beam type | Split |
| Pulse duration (ms) | 0.256 |
| Transmit Power (W) | 100 |
| Transducer gain (dB) | 18.00 |
| Major beam angle (degrees) | 17.1 |
| Minor beam angle (degrees) | 17.1 |
| Data Analysis Parameters |  |
| $S_v$ analysis threshold (dB) | -54 |
| TS analysis threshold (dB) | -48 |
| Elementary distance sampling unit (m) | 100 |
| Vertical bins (m) | 1 |
| Fish track detection |  |
| Minimum no. pings | 1 |
| Minimum no. single targets | 1 |
| Maximum gap between single targets (pings) | 5 |

#### A2. Laboratory analysis

In the laboratory, phytoplankton samples were size fractioned by pouring 100 ml of the well-mixed 500 ml phytoplankton field samples through 295, 64, and 5 μm sieves. The particles collected on the 295 μm sieve were discarded while the particles collected on the 64 and 5 μm sieves were each diluted with 15 ml of distilled water. This was repeated three times to produce three samples for the 64–295 μm and 5–64 μm size fractions. The 64–295 μm (large) phytoplankton were sized and counted using the FlowCAM 8000 (Yokogawa Fluid Imaging Technologies, Inc.) with a 4X objective and the 300 μm flow cell. The 5–64 μm (small) phytoplankton were sized and counted using a 10X objective and the 80 μm flow cell. Approximately 0.5–1 ml and 2–3 ml of the 15 ml samples was imaged, sized, and counted for the large and small phytoplankton samples, respectively. Two or three replicates for the phytoplankton sizes and abundances were processed for the six lakes.

Sub-samples (300 ml) of the zooplankton field samples were filtered through a 2000 μm sieve. Particles greater than 2000 μm were discarded. The sample was then diluted to 3 L and zooplankton (250–2000 μm in size) were counted and sized using the FlowCam Macro (Yokogawa Fluid Imaging Technologies, Inc.) with the 0.5X objective and the 2000 μm flow cell. The same 3 L sample was processed three times to produce replicates of the zooplankton sizes and abundances for each lake.

Non-planktonic particles such as bubbles, pollen, and fibers were removed from the images and size and abundance data using VisualSpreadsheet (Yokogawa Fluid Imaging Technologies, Inc.). FlowCAMs produce different estimates of size but the Area Based Diameter (ABD) estimate are similar to traditional microscopic body size measurements (Kydd et al., 2018). Therefore, the ABDs of the phytoplankton and zooplankton were used as the standard measure of body size.

### Supplementary Material B: comparing gillnet selectivity models

The gillnet data consist of individual fish size measurements of both fork length (*FL*, mm) and weight (*W*, g), caught by two different gillnet standards, the North American (NA) and the Ontario (ON), which represent the two groups affecting the observed size distributions, i.e., *g* = {NA, ON}. The gillnet size selectivity is a function of multiple mesh sizes *m_g,j_* (mm). The NA gear is composed of eight panels (*j* = 1, 2, …, 8) with mesh sizes ***m**_NA_* = {38,51, 64, 76, 89,102,114,127}; the ON gear is composed of five panels with mesh sizes ***m**_ON_* = {13,19,25,32,38}. The selectivity function of body size (*y*, fork length or weight) is represented by *s_g,j_*(*y, m_g,j_, **θ**_g_*) (***θ**_g_* represents the set of parameters determining the function shape) and requires paired (size, mesh) information for parameter estimation. Here we separate selectivity into two components: (i) a standardized function *r_g,j_*(*y, m_g,j_*) (***θ**_g_* has been omitted hereafter for simplicity) that varies from 0 to 1, which is useful for estimation purposes, and (ii) a gear-specific catchability coefficient *q_g_*, which determines the absolute selective value, so that *s_g,j_*(*y, m_g,j_*) = *r_g,j_*(*y, m_g,j_*)*q_g_*. Ideally, *r_g,j_*(*y, m_g,j_*) should include all size-dependent processes affecting selectivity, namely net encounter rates, contact rates (given encounter), and retention rates (given contact) (Anderson, 1998). However, the available data do not include marked populations and only allow the estimation of relative retention rates conditional on contact (Millar and Holst, 1997; Millar and Fryer, 1999). For this reason, we should interpret *r_g,j_*(*y, m_g,j_*) as a retention selectivity function. Given that encounter/contact rates can be approximated as a general power function of body size (Rudstam et al., 1984), ignoring them is expected to affect only the absolute estimates of size spectrum slope by a constant amount, not their relative variation across lakes. The lack of mark-recapture data, or any other information on absolute population sizes, also prevented the estimation of the absolute catchability coefficient *q_g_*. It could be estimated in relative terms only, by comparing the catch per unit effort from the mesh size common to both gear types (38mm). The following analysis will focus on comparing different models for retention selectivity *r_g,j_*(*y, m_g,j_*). The selected model was later used to get an expression for the probability density of observed sizes in gillnet catches (see Supplementary Material C).

There are three novel aspects of the present selectivity analysis worth mentioning:

1. it includes a modified size metric, here called *Effective Length (EL*), to account for differences in body shape between species;
2. it proposes a new function that better captures the observed shape of *r_g,j_*(*y, m_g,j_*);
3. estimation can be made with individual fish data points, so no arbitrary size binning is required.

Two fish with the same fork length *FL* but from different species can have markedly different vulnerabilities of capture by a given mesh, in a great part due to differences in body shape. One way to account for these differences is to use information on body weight to derive a linear size metric that standardizes for body shape across species. The simplest way is to impose an isometric, cubic relationship between length and weight. The resulting Effective Length (*EL*) is then given by:

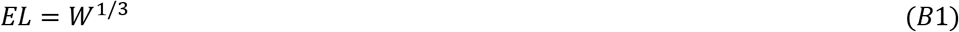

The models for gillnet retention selectivity are usually based on two main assumptions: (i) for each mesh size *m_g,j_* the selectivity has a maximum value (arbitrarily set to 1) at an intermediate body size 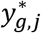, declining for smaller or larger sizes, and (ii) from the principle of geometric similarity, the shape and position of the retention curve 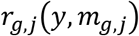 depends on the ratio *y/m_g,j_*, so both 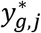 and the rate of decline in selectivity away from 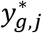 (i.e., the curve’s spread) are a fixed proportion of *m_g,j_*. For instance, two of the most commonly used functions are the normal and the log-normal (Millar and Fryer, 1999; Walker et al., 2013; Smith et al., 2017). The first predicts that the logarithm of *r_g,j_*(*y, m_g,j_*) will decline away from its maximum symmetrically as a quadratic function of the difference between *y/m_g,j_* and 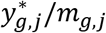; the second as a quadratic function of 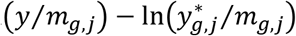. A preliminary assessment of these two alternatives can be made visually by plotting the relative catch or frequencies in a log-scale against ln(*y/m_g,j_*). This is done for the BsM data in Figure (B1), using the two alternative size metrics (*FL* vs *EL*).

**Figure B1.**
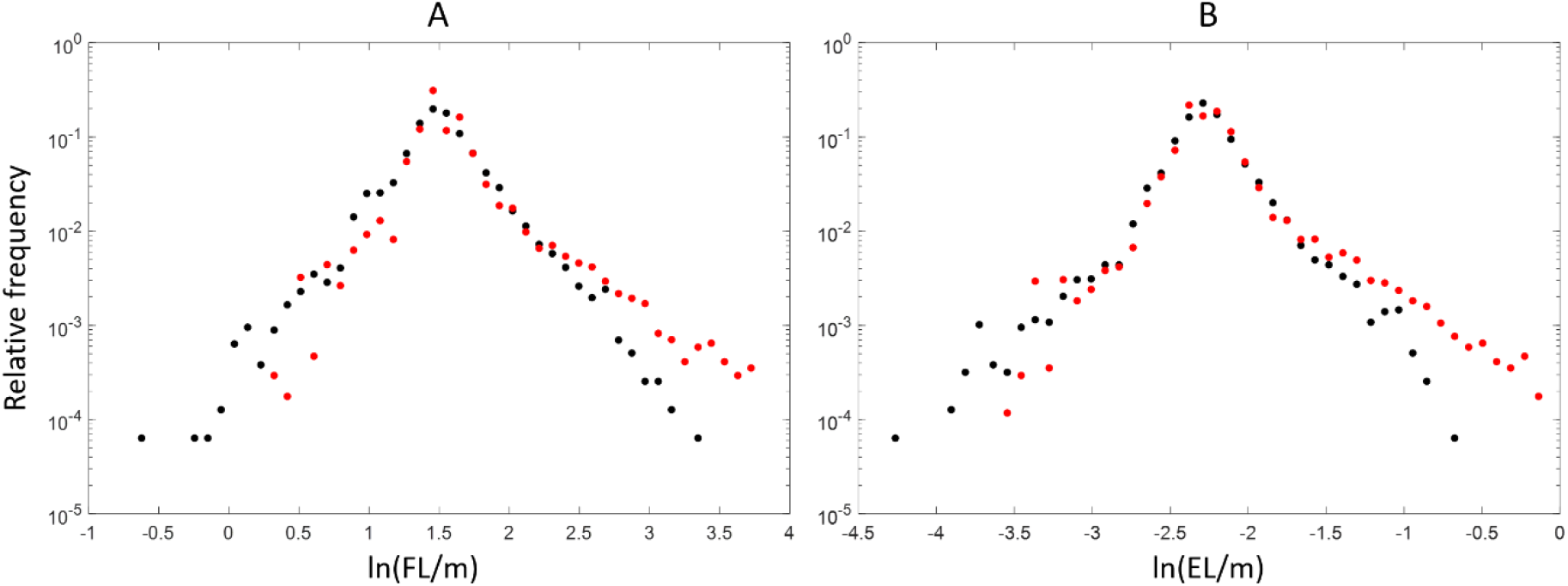
Distribution of NA (black dots) and ON (red dots) gillnet catches as a function of body size/mesh size ratio. *FL* = fork length (mm), *EL* = effective length (g^1/3^), *m* = mesh size (mm). Relative frequencies are plotted on a log-scale in all graphs.

The catch is approximately symmetrical when expressed as a function of ln(*y/m_j_*), which should favor the log-normal model. However, the pattern of selectivity decline away from its maximum does not follow a quadratic (parabolic) shape, it seems to be better approximated by a triangular shape instead. This suggests a symmetric exponential decline in vulnerability to catch as ln(*y/m_g,j_*) departs from its optimal value 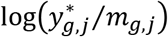. A model that satisfies this condition is given by Equation (B2):

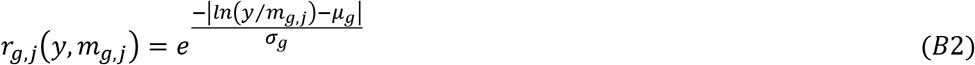

where 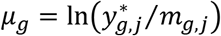 represents the body size/mesh size ratio (in a natural log-scale) with maximal vulnerability to catch (i.e., *r_g,j_*(*y, m_g,j_*) = 1) and *σ_g_* is a measure of how slowly the vulnerability declines as the log-ratio departs from *μ* (i.e., a measure of the curve’s spread). The total retention selectivity for a given size is the sum of Equation (B2) across mesh sizes, i.e., 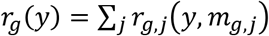. An example of this (hereby called) *Symmetric Exponential* selectivity curve is shown in Figure B2.

**Figure B2.**
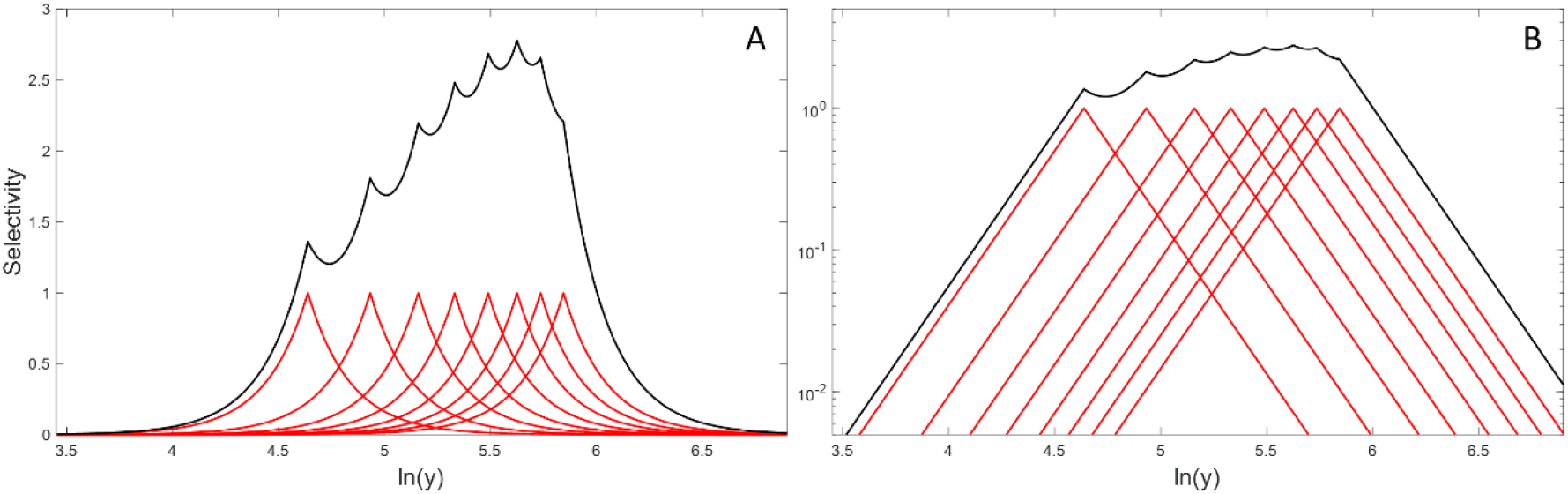
Symmetric Exponential retention selectivity curves *r_g,j_*(*y, m_g,j_*) (without tangling) plotted on a linear (A) and a log-scale (B) as a function of body size *y*. Red lines represent selectivity of individual gillnet mesh sizes *m_g,j_*; the black line is the total retention, summed across mesh sizes. Values of *m_g,j_* are the same as those for the NA gear (*j* = 1, 2, …, 8); the parameters were *μ_g_* = 1 and *σ_g_* = 0.2.

The estimation of selectivity parameters and model comparison was carried out through Bayesian inference. Twenty models were compared: five selectivity functions (Normal, Log-normal, Gamma, Inverse Gaussian, and Symmetric Exponential) (Millar and Fryer, 1999) x two size covariates (*FL* vs *EL*) x two tangling categories (presence vs absence of a tangling parameter). Adding tangling to a model has the effect of leveling off retention, leading to a constant *r_g,j_*(*y, m_g,j_*) value for *y/m_g,j_* ratios larger than a threshold. More specifically, the retention becomes *max*[*r_g,j_*(*y, m_g,j_*), *τ_g_*] for 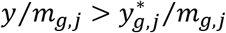, where the parameter *τ_g_* measures the rate of tangling (from 0 to 1) relative to maximum retention by the mesh. Walker et al. (2013) provide more background information on modelling the tangling component and examples of selectivity models with and without it.

The parameters of each model were assumed to be the same for NA or ON gillnets (e.g., ***θ**_NA_* = ***θ**_ON_* = {*μ, σ, τ*} for the Symmetric Exponential with tangling), so their selectivity curves were distinguished entirely due to their different mesh size compositions. This was necessary because estimation of some models failed to converge when parameters were made gear specific, which was probably due to low ON data resolution at small sizes. Although the distribution of body size/mesh size ratios is not identical for the two gears (Figure B1), using the same function for both is a reasonable approximation as the differences are mostly concentrated on the least frequent size classes (i.e., the right tail of the distributions).

The likelihood of observed data was calculated based on a variant of the SELECT method (Millar, 1992). Instead of using catch counts per mesh and size bin, here the basic data unit is the mesh *j* associated with each individual fish *i* caught. This is equivalent to assuming a multinomial distribution as is done with SELECT, but in this case based on a single event (the individual catch). The probability that fish *i* is caught by mesh *j*, given it has been caught by gear *g*, was:

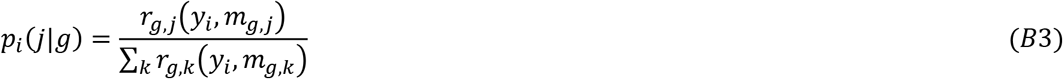

The total likelihood is the product of Equation (B3) across all individuals with mesh information, so the log-likelihood associated with the retention selectivity parameter set ***θ*** is expressed as:

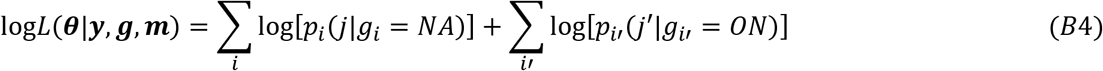

where *i* and *j* represent individuals and mesh sizes from NA nets, whereas *i*′ and *j*′ are individuals and mesh sizes from ON nets.

To estimate the selectivity parameters ***θ*** using Bayesian methods, we sampled their joint posterior distribution using the slice sampling MCMC algorithm in MATLAB R2020b. The log-posterior distribution is the sum of the log-likelihood function (Equation B4) with the log of prior distributions of ***θ***. The prior distributions were all non-informative and uniform. The range was [0,1000] for all positive parameters (i.e., k1 and k2 for the Normal and Inverse Gaussian, α and k for the Gamma model, *σ* for the Log-normal and Symmetric Exponential models) and [-1000,1000] for *μ* (Log-normal and Symmetric Exponential). Three independent chains were run for each model combination, each chain with burn-in of 1000, thinning of 10, and a final sample size of 1000. All chains converged to a common distribution, as indicated by the Gelman-Rubin statistic values (all < 1.1, Gelman and Rubin, 1992). Model selection was based on the Deviance Information Criterion (DIC, Spiegelhalter et al., 2002).

The best model (lowest DIC) was the Symmetric Exponential function with effective length (*EL*) as a size covariate and a tangling parameter (Table B1). Each one of these factors contributed independently for a better model fit, i.e., the Symmetric Exponential was the best model for any combination of size metric and tangling, *EL* was the best metric for any combination of selectivity function and tangling, and the presence of tangling was the best for any combination of selectivity function and size metric. The second-best model was the Normal with *EL* and tangling. Without the tangling parameter, the Normal function figures as the worst model, which suggests the addition of tangling partially compensates for the inability of the Normal to represent the markedly asymmetric (on a linear scale) distribution of body size/mesh size ratios. Figures B3–B7 show observed and predicted catches for all models with tangling. The size variables (x-axis) were binned for graphical purposes only (i.e., binning is not required for estimation).

**Table B1.** Deviance Information Criterion (DIC) and ΔDIC for the twenty combinations of selectivity function with or without a tangling parameter (rows) and size metric (columns).

| Function | Tangling | DIC $\times 10^{-4}$ | | ΔDIC $\times 10^{-4}$ | |
| --- | --- | --- | --- | --- | --- |
|  |  | Total length | Effective length | Total length | Effective length |
| Sym. Exponential | yes | 7.847 | 7.394 | 0.453 | 0 |
|  | no | 7.989 | 7.545 | 0.595 | 0.151 |
| Normal | yes | 7.988 | 7.506 | 0.594 | 0.112 |
|  | no | 10.714 | 10.659 | 3.320 | 3.265 |
| Log-normal | yes | 8.162 | 7.718 | 0.768 | 0.324 |
|  | no | 9.090 | 8.763 | 1.696 | 1.369 |
| Gamma | yes | 8.286 | 7.872 | 0.892 | 0.478 |
|  | no | 8.698 | 8.359 | 1.304 | 0.964 |
| Inv. Gaussian | yes | 8.360 | 7.956 | 0.966 | 0.562 |
|  | no | 8.782 | 8.460 | 1.388 | 1.066 |

**Figure B3.**
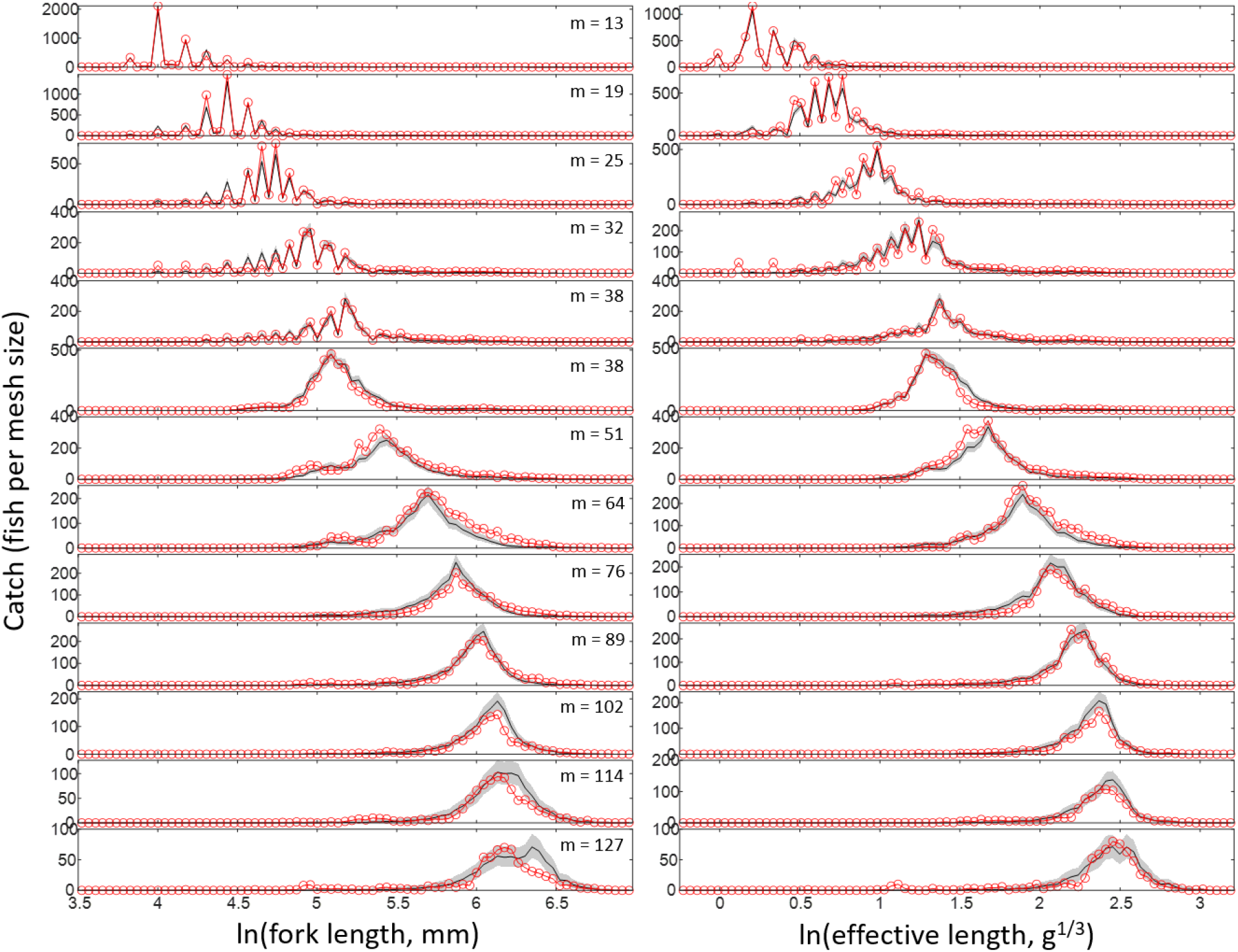
Observed (red lines) and predicted (gray bands = 95% prediction intervals, black line = median) catch for the **Symmetric Exponential** model with tangling and two size metrics (fork length and effective length). The top five mesh size panels are from ON gillnets, the bottom eight panels are from NA gillnets. In each panel, body size was divided into equally spaced bins on a log-scale. Predictions were generated based on the bin mid-point value, and 10 Multinomially distributed replicate values were generated for each one of the 3000 Bayesian samples. The expected value used for the Multinomial distribution was given by the product of *p*(*j*) (Equation B3) and the total catch for the size bin.

**Figure B4.**
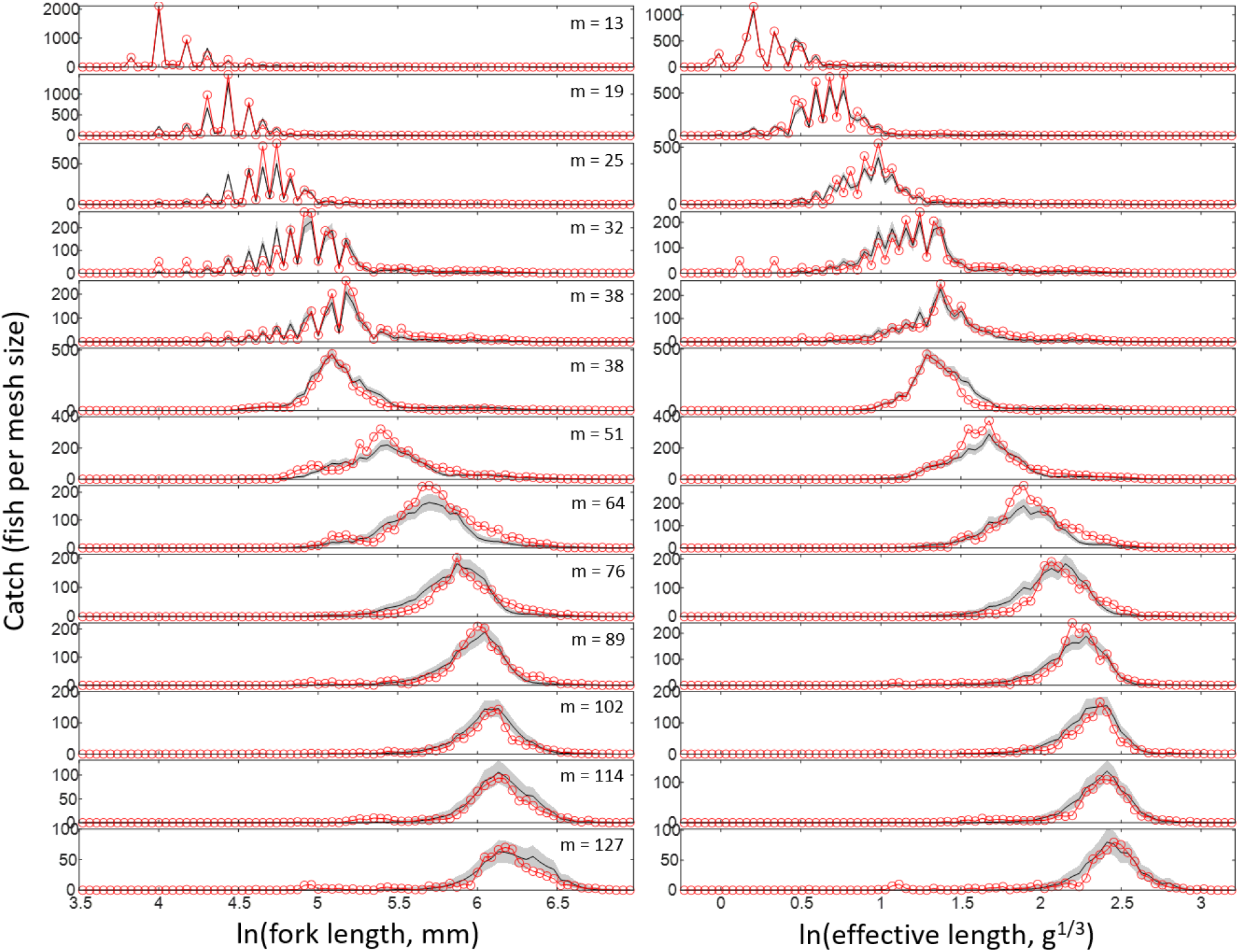
Observed (red lines) and predicted (gray bands = 95% prediction intervals, black line = median) catch for the **Normal** model with tangling and two size metrics (fork length and effective length). The top five mesh size panels are from ON gillnets, the bottom eight panels are from NA gillnets. In each panel, body size was divided into equally spaced bins on a log-scale. Predictions were generated based on the bin mid-point value, and 10 Multinomially distributed replicate values were generated for each one of the 3000 Bayesian samples. The expected value used for the Multinomial distribution was given by the product of *p*(*j*) (Equation B3) and the total catch for the size bin.

**Figure B5.**
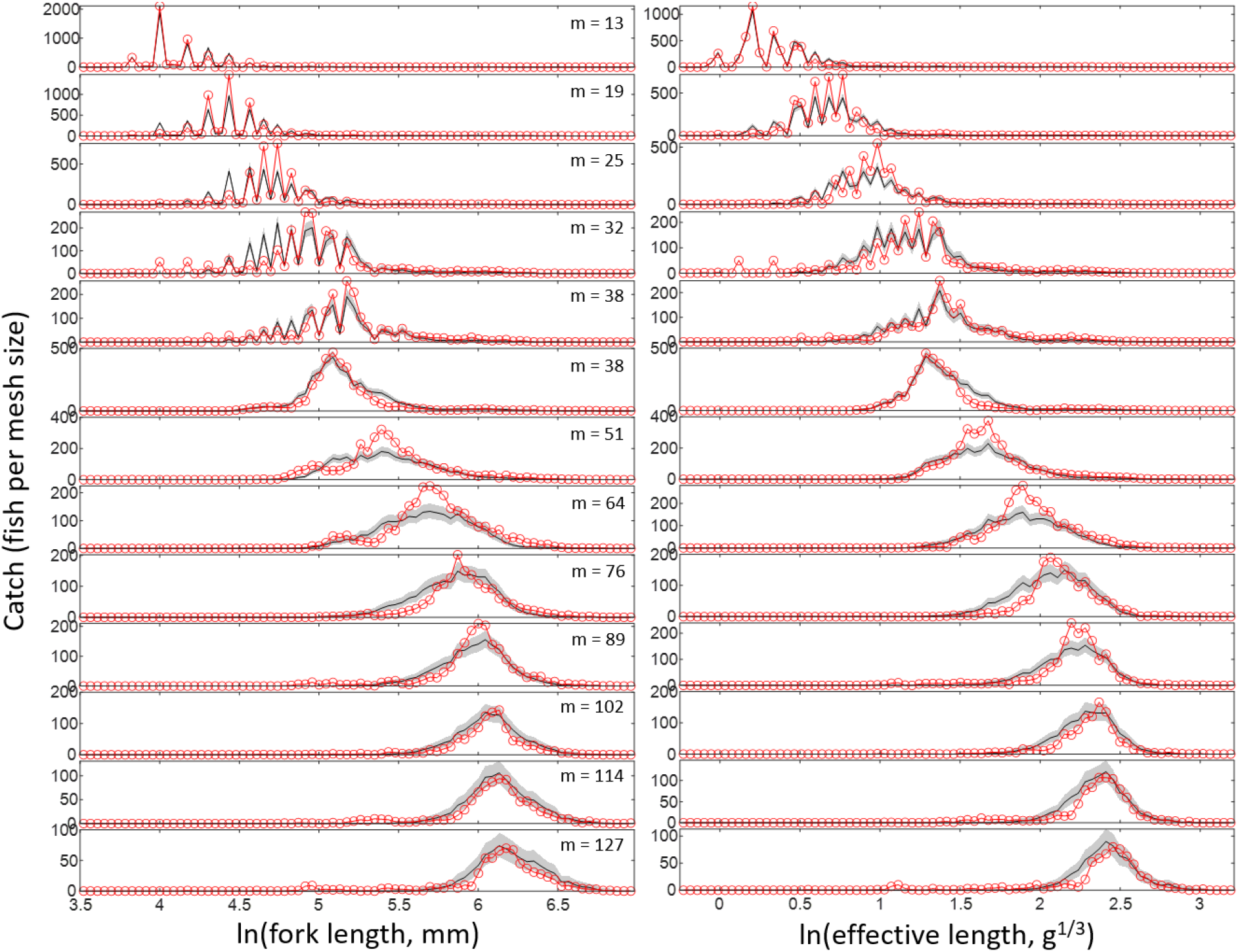
Observed (red lines) and predicted (gray bands = 95% prediction intervals, black line = median) catch for the **Log-normal** model with tangling and two size metrics (fork length and effective length). The top five mesh size panels are from ON gillnets, the bottom eight panels are from NA gillnets. In each panel, body size was divided into equally spaced bins on a log-scale. Predictions were generated based on the bin mid-point value, and 10 Multinomially distributed replicate values were generated for each one of the 3000 Bayesian samples. The expected value used for the Multinomial distribution was given by the product of *p*(*j*) (Equation B3) and the total catch for the size bin.

**Figure B6.**
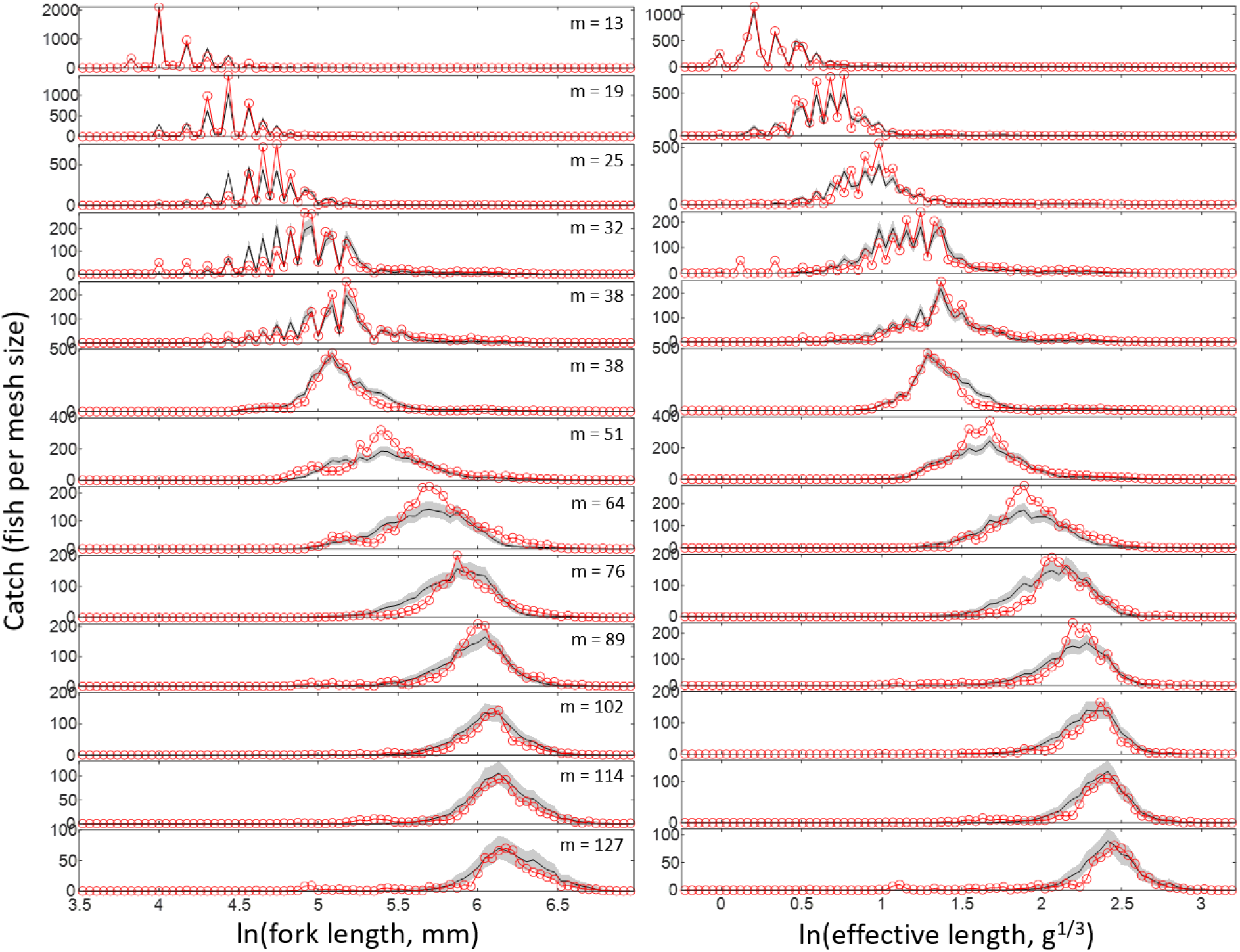
Observed (red lines) and predicted (gray bands = 95% prediction intervals, black line = median) catch for the **Gamma** model with tangling and two size metrics (fork length and effective length). The top five mesh size panels are from ON gillnets, the bottom eight panels are from NA gillnets. In each panel, body size was divided into equally spaced bins on a log-scale. Predictions were generated based on the bin mid-point value, and 10 Multinomially distributed replicate values were generated for each one of the 3000 Bayesian samples. The expected value used for the Multinomial distribution was given by the product of *p*(*j*) (Equation B3) and the total catch for the size bin.

**Figure B7.**
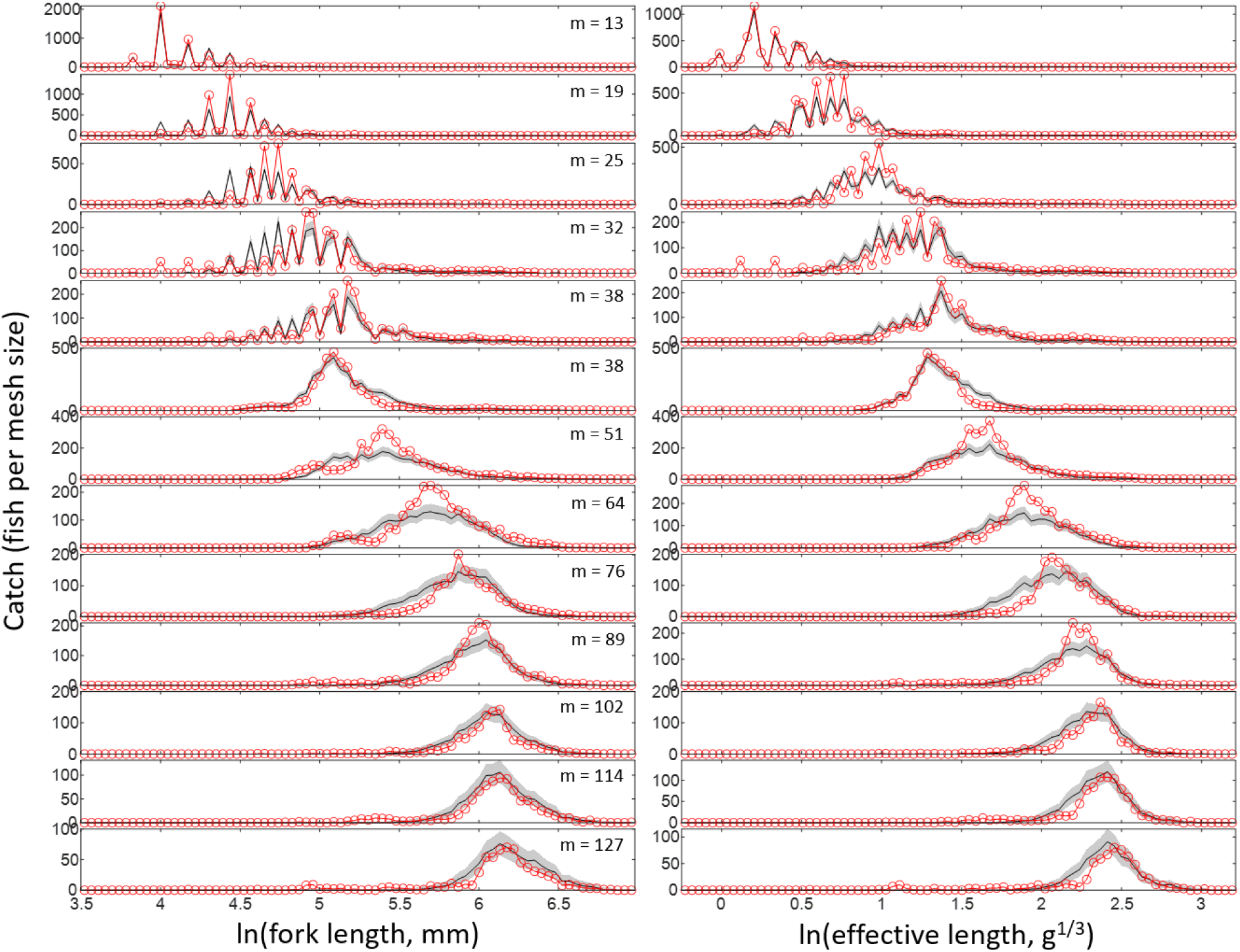
Observed (red lines) and predicted (gray bands = 95% prediction intervals, black line = median) catch for the **Inverse Gaussian** model with tangling and two size metrics (fork length and effective length). The top five mesh size panels are from ON gillnets, the bottom eight panels are from NA gillnets. In each panel, body size was divided into equally spaced bins on a log-scale. Predictions were generated based on the bin mid-point value, and 10 Multinomially distributed replicate values were generated for each one of the 3000 Bayesian samples. The expected value used for the Multinomial distribution was given by the product of *p*(*j*) (Equation B3) and the total catch for the size bin.

### Supplementary Material C: probability distributions and likelihoods

#### C1. General model

The size spectrum describes the decline in abundance (*n*, in number of organisms per habitat area or volume, per unit of body size) with body size (*y*, weight or length) according to a power law: *n* = *κy*^−*λ*^. The probability density function of *y* is proportional to *n*, but normalized so that it integrates to 1 within its size domain (*y_min_*≤*y* ≤ *y_max_*):

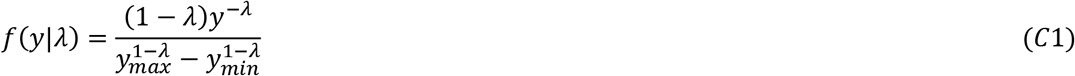

where *f*(*y*|*λ*) is the probability density function of *y* conditional on its parameter and *λ*, which is the magnitude of the size spectrum slope (the slope itself is −*λ*). As pointed out in the main text, the term *slope* has been traditionally used in size spectrum research, and for this reason has been adopted here, despite −*λ* being an exponent in Equation (C1). Although the real size distributions may not strictly follow Equation (C1), the function gives a theoretical description of their primary structure, i.e., the first-order decline in abundance from small to large organisms, whose indicator *λ* can be linked to ecosystem processes related to individual metabolism, size-dependent predation, and trophic transfer efficiency (Andersen and Beyer, 2006; Trebilco et al., 2013; Mehner et al., 2018). We expect that, by finding the estimate of *λ* that best describes the observed size distribution of a given community, we will be giving a reliable indicator of its primary structure and therefore of those ecosystem processes.

The general approach for estimating *λ* is to derive another probability density function for body size that combines (i) the underlying size spectrum *f*(*y*|*λ*) with (ii) a group- or gear-specific selectivity function *s_g_*(*y, **θ**_g_*), which measures the probability of detecting or catching any individual with size *y*, per unit sampling effort, conditional on the selectivity parameter set ***θ**_g_* (which is composed by multiple parameters that determine the shape and magnitude of *s_g_*) and (iii) the sampling effort associated with the group or gear, *v_g_*, measured as e.g., water volume (m^3^) in hydroacoustic surveys or total length of nets (m) in gillnet surveys. The resulting function, denoted here by *o_g_*(*y*|*λ, **θ**_g_*), gives the expected probability density of observed body sizes in a sample of group *g* (or collected by gear *g*). The relative probability of observing a given *y* is simply the product of its underlying probability density in the wild, i.e., *f*(*y*|*λ*), and the probability that it is caught in a sample with effort *v_g_*, i.e., *s_g_*(*y, **θ**_g_*)*v_g_*. To derive *o_g_*(*y*|*λ, **θ**_g_*), an additional necessary step is to normalize the resulting product so that the total probability of observing an individual of any size integrates to 1:

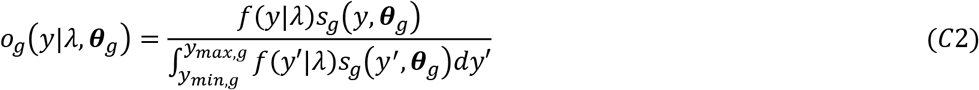

where the normalization constant in the denominator is the product’s integral over the group-specific size domain from *y_min,g_* to *y_max,g_*. Notice that in this group-specific case the sampling effort *v_g_* cancels out for being both in the numerator and denominator, so it is absent in Equation (C2). If only one group is part of the sample, then Equation (C2) would suffice to calculate the likelihood associated with *λ* for each individual observation *i*, i.e., *L*(*λ*|***θ**_g_, y_i_*) = *o_g_*(*y_i_*|*λ, **θ**_g_*). With the likelihood function and a prior distribution for *λ*, Bayesian methods can then be applied to estimate the posterior distribution conditional on the observed data and known selectivity parameters, i.e., *p*(*λ*|***θ**_g_,**y***), which provides all desired estimates such as the mean (or median) *λ* and its uncertainty (e.g., standard deviation, interquartile range, or 95% credible interval). If the selectivity parameters are not yet known but are jointly estimated with *λ*, then another likelihood function will be necessary, in this case associated with ***θ**_g_* and additional gear-specific data ***X*** (such as a matrix with individual size, gear and mesh information from gillnets), i.e., *L*(***θ**_g_*|***X***). With both likelihood functions and priors for *λ* and all ***θ**_g_*, a joint posterior *p*(*λ, **θ**_g_*|***y,X***) can be estimated considering uncertainty in both the size spectrum and the gear selectivity parameters.

In case the survey data include two or more groups (or gear types), the observed size distribution in a sample will depend explicitly on the relative sampling intensities or efforts ***v*** = {*v_g_*; *g* = 1,2,..}. The resulting distribution *o_g_*(*y*|*λ, **θ, v***) will be a mixture distribution, i.e., the sum of the individual group distributions (*o_g_*(*y*|*λ, **θ**_g_*), Equation C2) weighted by the probability that any given observation belongs to that group, *ρ*(*g*|*λ, **θ, v***), i.e.,

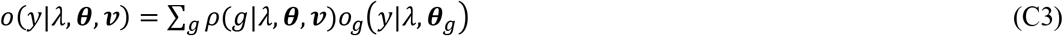

The probability *ρ*(*g*|*λ, **θ, v***) is equal to the total expected number of individuals of group *g* in the sample, relative to the total from all groups, and is dependent on the size spectrum and selectivity parameters:

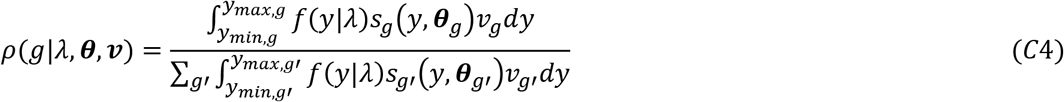

which ensures that ∑*_g_ ρ*(*g*|*λ, **θ,v***) = 1. The resulting total distribution after combining Equations (C2–C4) is:

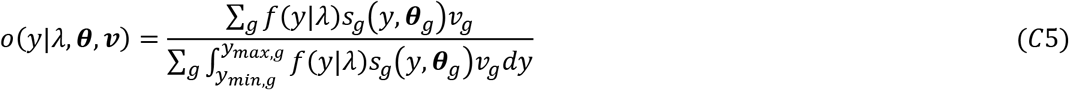

In usual size-spectrum surveys, each individual observation *i* can be assigned to both a group category *g_i_* and a body size *y_i_*. In this case, the individual likelihood will be equal to the probability of belonging to group *g* multiplied by the probability density of *y* conditional on that group, i.e., *L*(*λ*|***θ**, y_i_, g_i_, **v***) = *ρ*(*g_i_*|*λ, **θ, v***)*o_g_*(*y_i_*|*λ, **θ**_g_*). The total likelihood is the product of all individual likelihoods:

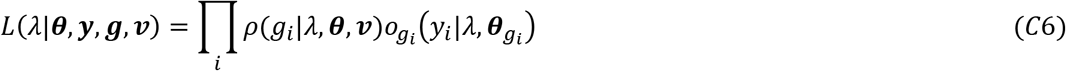

and the log-likelihood is:

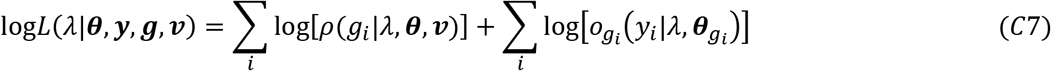

As part of notation for model development, we use *log* to represent a generic logarithmic function, which is applicable to any logarithmic basis as long as the basis is used consistently within the same analysis. Whenever a single basis was used for specific purposes (e.g., natural logarithm, log_2_ or log_10_), we indicate it explicitly.

The first summation term of Equation (C7) gives the likelihood due to group identities (based on Equation C4) and the second term gives the likelihood due to the observed size distribution within a group (based on Equation C2). In case there is no individual size information, but only the total number of observations and sampling effort for each group (or sampling gear), the log-likelihood simplifies to the first summation term:

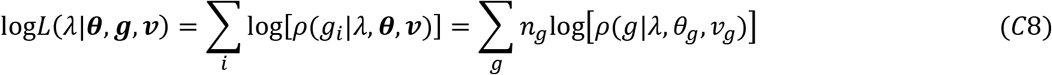

where *n_g_* is the total number of individuals from group *g*. If, alternatively, we ignore information on group-specific sampling efforts, the log-likelihood becomes:

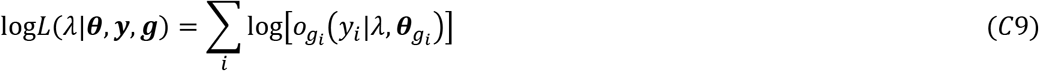

Still another possibility is if we know individual sizes and sampling efforts but have no information on the group each individual belongs to. In this case, the log-likelihood will be based on the mixture distribution (Equation C5):

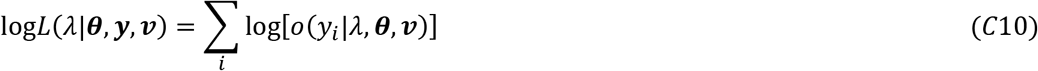

#### C2. Community data

The community data was structured as counts of individuals per size bin *b*, based on body length (mm), whose lower limits are given by *y_b_*. The limits of size bins were logarithmically spaced on a basis 2 with fixed width = 0.25, i.e., log_2_(*y*_*b*+1_) – *log*_2_(*y_b_*) = 0.25. Each major trophic group *g* was defined by a size interval from *y_min,g_* (the lower limit of the group’s first bin) to *y_max,g_* (the upper limit of the group’s last bin). For phytoplankton (*g* =1): *y*_*min*,1_ = 2^−7.875^, *y*_*max*,1_ = 2^−1.875^; for zooplankton (*g* =2): *y*_*min*,2_ = 2^−1.875^, *y*_*max*,2_ = 2^1.125^; for fish (*g* =3): *y*_*min*,3_ = 2^5.875^, *y*_*max*,3_ = 2^10.375^. The minimum for each group was defined so that all lakes have non-zero counts for the first group’s bin, and the maximum was defined so that at least one lake have non-zero counts in the last group’s bin. This criterion also ensured no size overlap between trophic groups.

These size limits define the domain of the community’s observable size distribution. In other words, the probability of observing an individual outside those bounds for each group was set to zero. These bounds reflect limitations of sampling methods and detectability rather than actual boundaries in the distribution of organisms in the community, whose range should be broader. Within the observable limits, organisms were assumed to be equally catchable and detectable by the sampling gear, so the selectivity function defining the probability of detection or capture is given by:

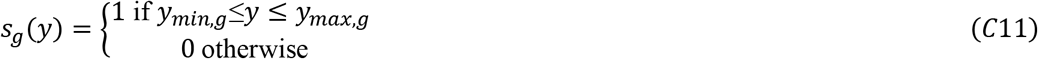

The observed size distribution within a given group’s size range, *o_g_*(*y*), is the product between the underlying size spectrum (Equation C1) and the selectivity function, normalized so that it integrates to 1 (Equation C2). To get the probability that an individual belongs to size bin *b*, we integrate *o_g_*(*y*) within the bin limits to get an expression for the observed distribution of size bins:

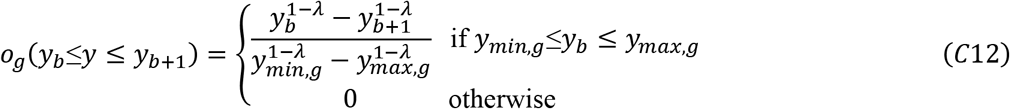

Different from the general case in the previous section, *o_g_*(*y_b_*|*y* ≤ *y*_*b*+1_) is a discrete probability distribution whose value is the actual probability of observing an individual in size bin *b*, given it belongs to group *g*. To get the probability that an individual belongs to group *g*(*ρ*(*g*), Equation C4), we multiply the total expected number of individuals in that group, i.e., 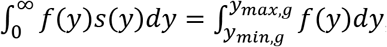 by the sampling effort associated with that group, *v_g_*, which in this case is the sampled volume (m^3^). This leads to:

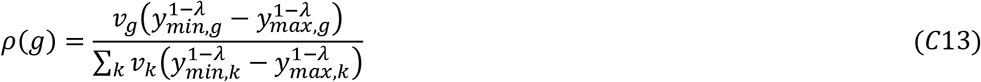

As the size intervals characterizing groups do not overlap in our case, the final probability associated with bin *b* will be simply the product *ρ*(*g_b_*)*o_g_b__*(*y_b_*≤*y* ≤ *y*_*b*+1_) for the group containing that bin, *g_b_*. The observed data are structured as counts per size bin, represented by the vector ***c*** = [*c*_1_, *c*_2_,…, *c_B_*], with a total count 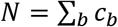. The likelihood of observing ***c*** in a survey is thus defined by a multinomial distribution:

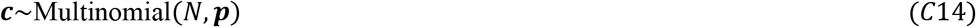

where the parameter vector ***p*** contains the bin probabilities *p_b_* = *ρ*(*g_b_*)*o_g_b__*(*y_b_*≤*y* ≤ *y*_*b*+1_). This is similar to the binned method of Edwards et al. (2017, 2020) for deriving likelihoods, but here it combines multiple groups to get a single spectrum estimate.

The log-likelihood function of the multinomial distribution, after excluding terms that do not involve the parameter of interest (*λ*), can be separated into its *ρ* and *o_g_* components:

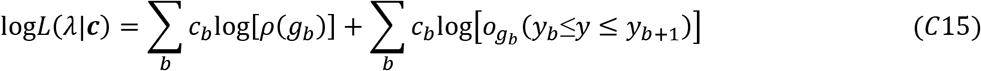

which can be linked to *λ* through Equations (C12–C13). To estimate *λ* based on *between-group* variation (Equation C8), the log-likelihood function simplifies to the first summation term of Equation (C15), whereas for *within-group* variation (Equation C9), the log-likelihood function simplifies to the second summation term.

#### C3. Gillnet data

The gillnet data consisted in individual size measurements (fork length and weight) from two gears: the North American (NA, or large-mesh) and the Ontario (ON, or small-mesh) gillnet standards, both part of the Broad-scale Monitoring protocol (BsM, Sandstrom et al., 2013). Part of the ON data, particularly regarding the smallest mesh sizes, originated as individual counts within size bins of 1cm (based on fork length), which were later converted to individual size measurements using the bin mid-point and weight-length relationships to convert to weight values (Supplementary Material A). Although fully accounting for the original binning information is preferable in terms of accuracy (Edwards et al., 2020), our analysis relied on the individual size conversions for three reasons: (i) we wanted to illustrate the application of the method to samples of individual size measurements, which are common in gillnet monitoring surveys, (ii) most of the data were originally collected as individual measurements, including the entirety of the NA catches and 40% of the ON catches, (iii) it greatly simplifies the analysis. Fully accounting for the bin structure is feasible and would involve a hybrid approach combining the likelihood calculation methods of the previous section (C2. *Community data*) with the individual-based likelihoods of this section.

To further simplify analysis, it is convenient to represent the size variable on a log-scale, given that the chosen selectivity model is a function of ln(*y*) (Supplementary Material B). It is also convenient to use body weight (*W*) as our size metric, as the best covariate explaining the mesh-specific catches (Effective length, *EL*) is a direct function of *W*. Let’s thus define:

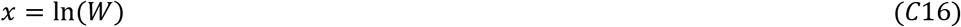

If we replace *y* by *W* in Equation (C1) and apply Equation (C16) (*W* = *e^x^*), the size spectrum of *x* becomes an exponential with a generic form *f*(*x*|*λ*) = *C*(*e^x^*)^−*λ*^ = *Ce*^−*λx*^, where *C* is a constant (independent of *x*) ensuring *f*(*x*|*λ*) integrates to 1 (i.e., that it is a proper probability density function) within its domain from *x_min_* to *x_max_*. By setting 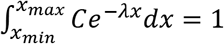, one finds that *C* = *λ*/[*e*^−*λx_min_*^ – *e*^−*λx_max_*^], and the resulting probability density function is:

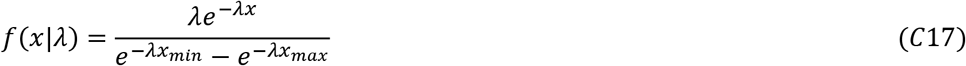

The distribution of observed sizes *o_g_*(*x*|*λ, **θ**_g_*) caught by gillnet gear *g* will be distorted by the gear selectivity *s_g,j_*(*EL*(*x*), *m_g,j_, **θ**_*g*_*), where *EL*(*x*) is the Effective Length (Equation B1), *m_g,j_* is the size of mesh *j* (mm) from gear *g*, and ***θ**_g_* is the set of parameters determining the shape of the selectivity curve. The total selectivity for a given *EL* is the sum of selectivity functions across mesh sizes, i.e., 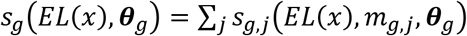. Combining Equation (B1) and (C16), the Effective Length becomes *EL*(*x*) = *e*^*x*/3^. Selectivity is assumed to be the product of a relative retention selectivity component *r_g,j_*(*e*^*x*/3^, *m_g,j_, **θ**_g_*) and an absolute catchability coefficient *q_g_* (Supplementary Material B) Therefore, *o_g_*(*x*|*λ, **θ**_g_*) is proportional to the product *f*(*x*|*λ*)*s_g,j_*(*EL*(*x*), *m_g,j_, **θ**_g_*) = *f*(*x*|*λ*)*r_g,j_*(*e*^*x*/3^, *m_g,j_, **θ**_g_*)*q_g_*. To estimate the size spectrum parameter *λ*, it is necessary that *o_g_*(*x*|*λ, **θ**_g_*) be a true probability density function, which is achieved by dividing the product by its definite integral over the size domain from its minimum (*x_min_*) to maximum (*x_max_*), i.e.:

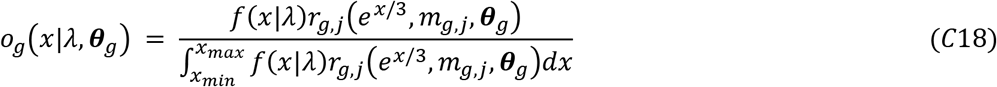

which is analogous to the general Equation (C2). When only one gear type (e.g., NA) is considered, the catchability coefficient *q_g_* cancels out (i.e., only relative catchabilities matter), not appearing on Equation (C18). Using the symmetric exponential model (Equation B2) for *r_g,j_*(*e*^*x*/3^, *m_g,j_, **θ**_g_*), the resulting probability density of observed sizes within gear *g* for mesh size *j* is:

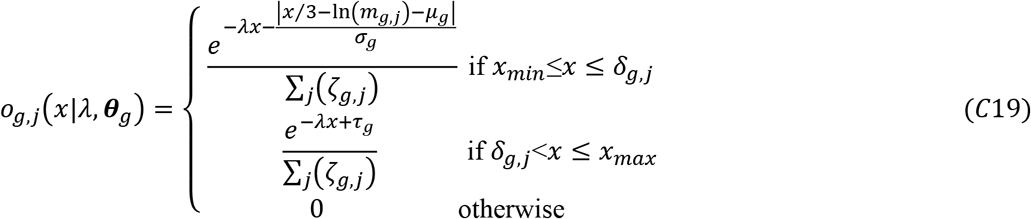

where 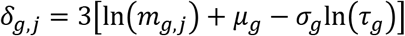 is the threshold size above which retention becomes equal to the tangling parameter *τ_g_* (Figure C1). To get the total probability density of *x* within gear *g*, we sum across mesh sizes:

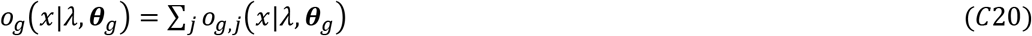

The term 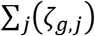 in the denominators of Equation (C19) is the integration constant ensuring that the total probability *o_g_* integrates to 1. The mesh-specific term *ζ_g,j_* is the sum of three components, i.e., *ζ_g,j_* = *ζ*_*g,j*,1_ + *ζ*_*g,j*,2_ + *ζ*_*g,j*,3_, where:

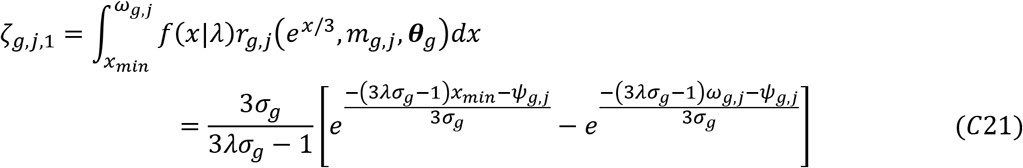

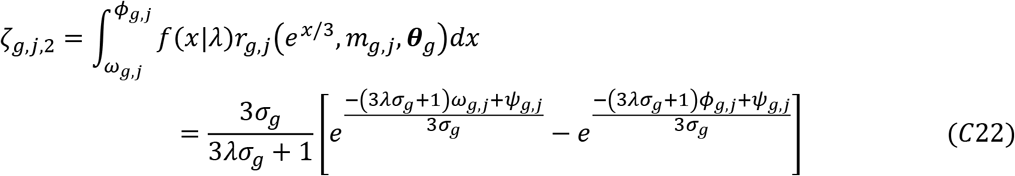

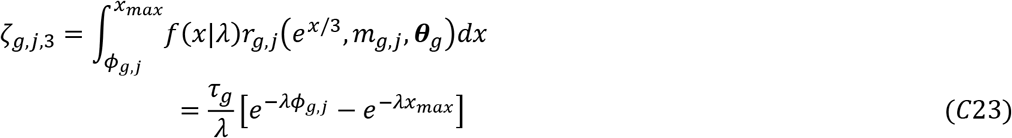

The first is the integral corresponding to the ascending part of the selectivity curve before it peaks at *x* = *ψ_g,j_* = 3[ln(*m_g,j_*) + *μ_g_*] (Figure C1); the second is the integral corresponding to the descending part after the peak, but before retention starts being determined by tangling at *x* = *δ_g,j_* = 3 [ln(*m_g,j_*) + *μ_g_* – *σ_g_*ln(*τ_g_*)]; the third is the integral defined after that tangling threshold. The terms *ω_g,j_* = max[*x_min_*, min(*x_max_, ψ_g,j_*)] and *φ_g,j_* = max[*ω_g,j_*,min(*x_max_, δ_g,j_*)] ensure that the integral is taken within the correct size interval and the appropriate part of the function (e.g., ascending or descending). For instance, if the maximum size *x_max_* is smaller than the size at peak, *ψ_g,j_*, then *ω_g,j_* = *φ_g,j_* = *x_max_*, as a consequence *ζ*_*g,j*,2_ = *ζ*_*g,j*,3_ = 0 and only the first, ascending part of the curve will count for the full integral (*ζ_g,j_* = *ζ*_*g,j*,1_). Examples of the probability function *o_g_*(*x*) are shown in Figure C2 for different values of *λ* and *g* = NA.

**Figure C1.**
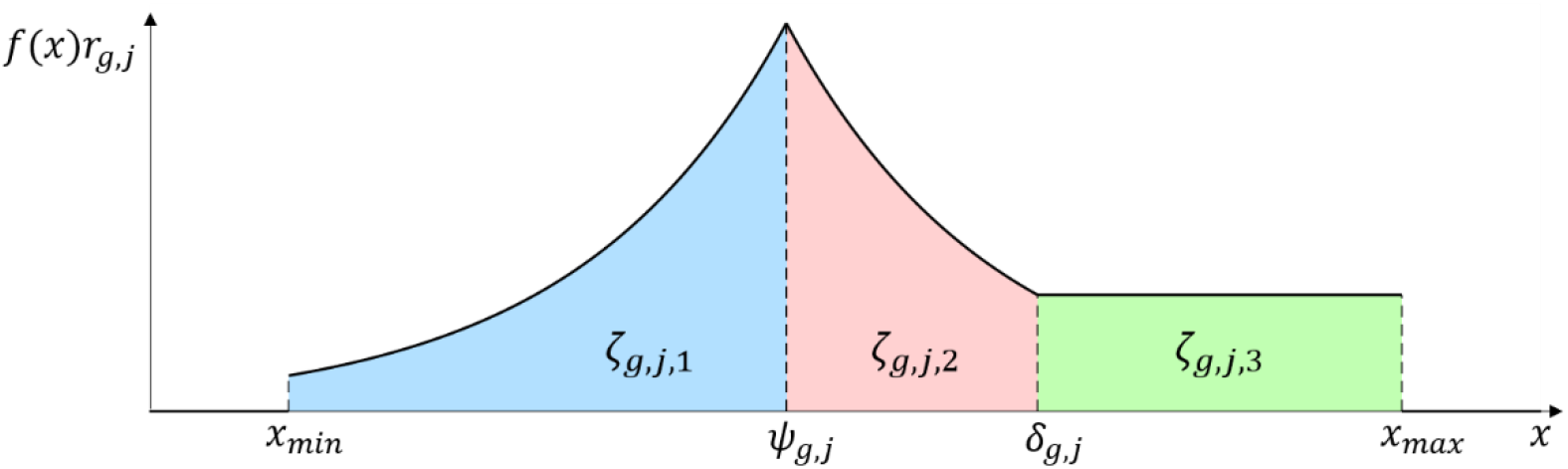
Illustration of the three integrals from Equations (C21–C23) (*ζ*_*g,j*,1_, *ζ*_*g,j*,2_, and *ζ*_*g,j*,3_, different colloured areas). The function *f*(*x*)*r_g,j_* (thick black line) is the product of the size spectrum and the retention selectivity function (here the spectrum parameter *λ* ≈ 0, so the function has the same shape as *r_g,j_*). The body size (*x*) reference points, marked by vertical dashed lines, are: minimum size (*x_min_*), maximum size (*x_max_*), size at the peak of retention (*ψ_g,j_*), size threshold for the tangling component (*δ_g,j_*) (i.e., the size above which *r_g,j_* = *τ_g_*).

**Figure C2.**
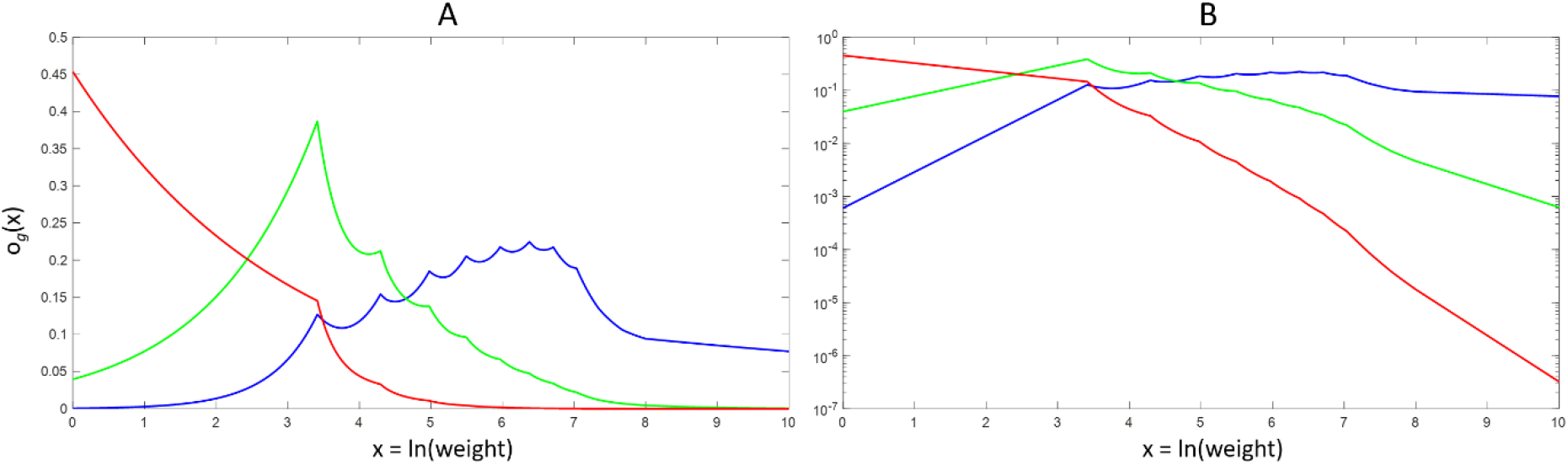
Probability distribution of observed sizes, *o_g_*(*x*) (Equation C20), plotted on a linear (A) and on a log scale (B) for three different values of size spectrum slope: *λ* = 0.1 (blue), 1 (green), and 2 (red). Mesh sizes were the same as those for the NA gear (*j* = 1, 2, …, 8); the selectivity parameters were *μ* = −2.5, *σ* = 0.2 and *τ* = 0.2.

The probability that an individual was caught by gear *g, ρ*(*g*|*λ, **θ, v***), is proportional to the integral of the gear’s function 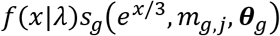, multiplied by the sampling effort *v_g_* (measured as total extension of a mesh panel multiplied by the number of net gangs, in meters) relative to the total from all gears combined (Equation C4):

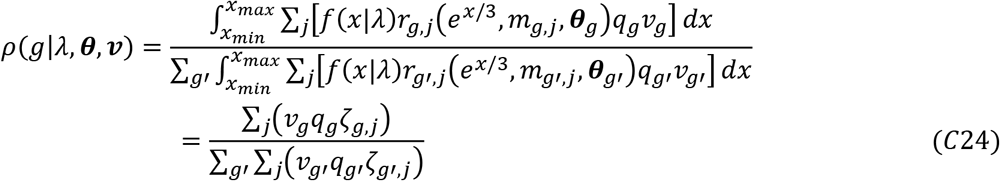

The probability density of *x* with both gears combined, *o*(*x*|*λ, **θ***) (Equation C5), is a mixture distribution resulting from the sum of *o_g_*(*x*|*λ, **θ**_g_*) across gears, weighted by their probabilities *ρ*(*g*|*λ, **θ, v***):

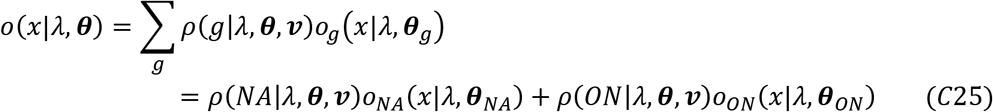

An example of this probability density is shown in Figure C3.

**Figure C2.**
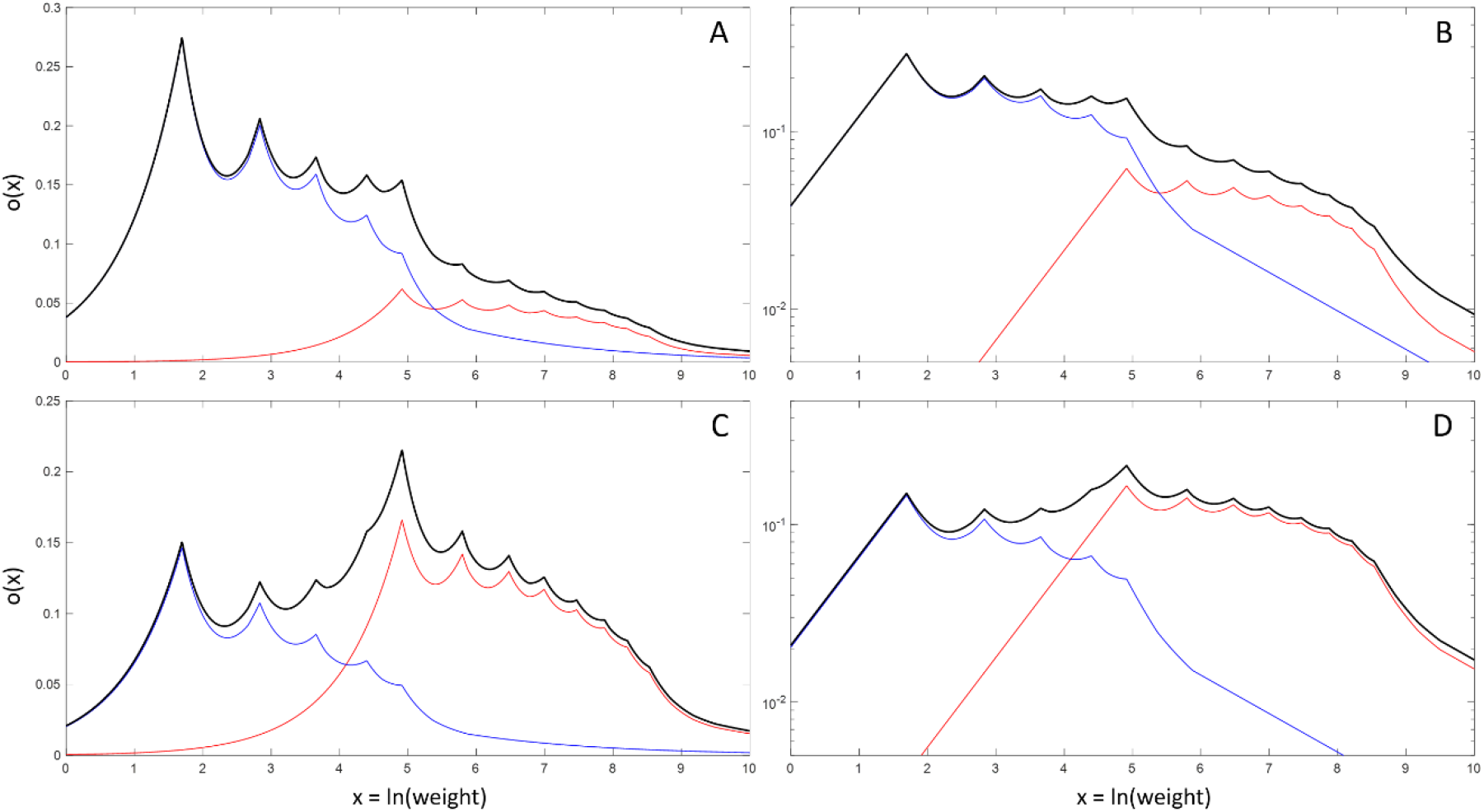
Probability distribution of observed sizes, *o*(*x*) (Equation C25, thick black lines), combining two gear types (plotted on a linear scale in A,C, and on a log-scale on B,D). The thin blue and red lines represent the ON and the NA gear, respectively, with their associated mesh size compositions (the plotted values are the product *ρ*(*g*)*o_g_*(*x*) for each gear). In (A-B), the effort x catchability product (*v_g_q_g_*) is the same for both gears; in (C-D), *v_NA_q_NA_* is five times greater than *v_ON_q_ON_*. Parameter values were: *λ* = 0.5, *μ* = −2, *σ* = 0.2 and *τ* = 0.2.

The likelihood of observing each individual fish with log-weight *x* and caught by gear *g*, *L*(*λ*|***θ**, x_i_, g_i_, **v***), is the product of Equations (C20) and (C24). The log-likelihood function associated with *λ* is the sum of log*L*(*λ*|***θ**, x_i_, g_i_, **v***) across all individuals caught (Equation C7). To account for uncertainty in the selectivity parameters and potential error propagation, we jointly estimated *λ* and ***θ***. This means that the likelihood of individual size and gear observations depended both on *λ* and ***θ***, i.e., *L*(*λ*|***θ**, x_i_, g_i_, **v***) ≡ *L*(*λ, **θ***|*x_i_, g_i_, **v***). The selectivity parameter set ***θ*** is composed by the Symmetric Exponential parameters *μ_g_*, *σ_g_*, and *τ*, assumed to be the same for both gears (Supplementary Material B), and the catchability coefficients *q_NA_* and *q_ON_*. Because absolute catchabilities are not known for either gear and there is no information allowing to estimate absolute population sizes from gillnet catches, the catchability coefficients could only be estimated in relative terms. For simplicity and without loss of generality, we imposed *q_NA_* = 1 and estimated *q_ON_* based on additional data from the BsM mesh-specific catches (explained below). The log-likelihood function associated individual fish catches from the study lakes, and based on Equation (C7), is:

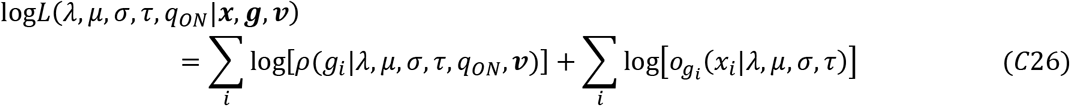

The total log-likelihood function included additional likelihood functions for the selectivity parameters based on mesh-specific information from the Broad-Scale Monitoring. The first of those, log*L*(*μ, σ, τ*|***x_M_, g_M_, m***), measures the likelihood that individual fishes with size *x* caught by gear *g* are found in mesh size *m_g,j_*, and is based on Equations (B3–B4) from Supplementary Material B:

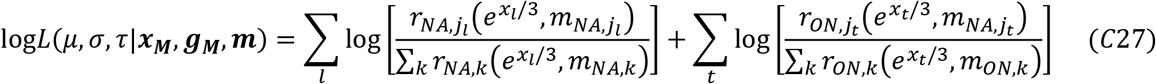

where ***x_M_*** and ***g_M_*** represent the vectors of log-weight values and gear identities from the mesh-specific BsM data (differing from the vectors ***x*** and ***g*** representing gillnet catches in the six study lakes), which are paired with the vector ***m*** containing mesh size information of individual catches; the subscripts *l* and *t* represent individuals caught by NA or ON nets, respectively, and 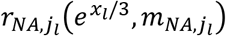 is the retention selectivity function (Equation B2, with tangling) for the NA gear, evaluated for individual *l* with Effective Length *EL* = *e*^*x_l_*/3^ and caught by mesh *j_l_* with size *m_NA,j_l__* (parameters *μ, σ, τ* were omitted from the function for simplicity).

The second likelihood function associated with mesh-specific data, log*L*(*q_ON_*|***g_38_, v_M_***), represents the chance that fishes caught by mesh size *m*=38 (common to both gears) belong to the ON gear. We assume that differences in catch-per-unit-effort from mesh 38 are representative of differences in catchability for the whole gears. Here the vector ***g*_38_** represents the gear identity of fishes caught by mesh size 38, and ***v_m_*** = {*v_M,NA_*, *v_M,ON_*} is the vector with the sampling efforts (total mesh panel extension, in meters) from the two gears that had information on mesh-specific catches. To control for differences in the spatial distribution of NA and ON nets, we considered as part of the effort and catch only the depth strata that had at least one of each gear. The probability that a fish from mesh 38 had been caught by an ON net is proportional to the product of its effort and catchability, relative to the total from both gears, i.e.:

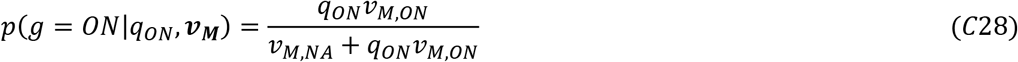

The total catch from mesh 38 belonging to ON nets, *c*_38,*ON*_, is thus expected to follow a binomial distribution with parameters *n* = *c*_38,*ON*_ + *c*_38,*NA*_ and *p* given by Equation (C28). The resulting log-likelihood function is:

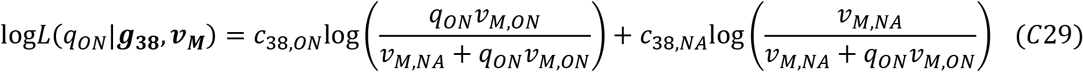

The total log-likelihood function is the sum of the above three likelihood functions:

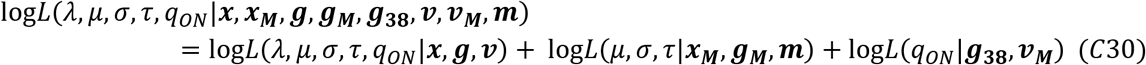

To estimate *λ* based on *between-gear* variation, we used Equation (C9) and (C24). We could not jointly estimate the selectivity parameters due to failure of convergence of the posterior distributions. These parameters were estimated separately through the second and third likelihood terms from Equation (C30), and their median values used as constants for estimation of *λ* through the log-likelihood function:

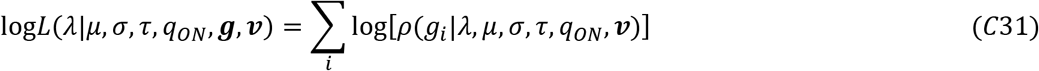

For the estimation of *λ* based on *within-gear* variation, we used Equation (C9) instead of (C7) for the first term of the total log-likelihood and ignored differences in catchability and effort. For consistency, the retention selectivity parameter values were the same as those used for the between-group estimation, so the log-likelihood function was:

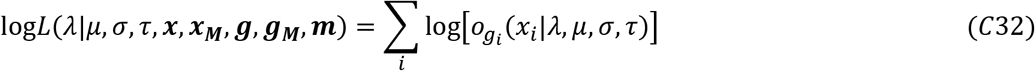

### Supplementary Material D: Bayesian analyses

The likelihood functions derived in Supplementary Material C provide the basic ingredients for parameter estimation, which can be carried out in two main ways (i) through Maximum Likelihood Estimation (MLE), in this case using numerical optimization methods (i.e., no closed analytical expression for the maximum likelihood could be obtained) or (ii) through Bayesian estimation, using Markov chain Monte Carlo (MCMC) methods (Stauffer, 2007). We opted for the later as it allows for a full characterization of the joint distribution of parameters (i.e., the posterior distribution). The posterior distribution gives the probability density of parameters given observed data and can be obtained through Bayes theorem as the product of two probabilities (which becomes a sum in a log-scale): (i) the probability of obtaining the observed data, given the parameters and (ii) the prior probability of parameters. The first is equivalent to the likelihoods derived in the Supplementary Material C, whereas the second must be informed as part of analysis using any available prior information about the parameters. For instance, the posterior distribution of parameters *λ, μ, σ, τ*, and *q_ON_* given all the observed gillnet data ***Y*** = {***x, x_M_, g, g_38_, g_38_, v, v_M_, m***}, i.e., *ρ*(*λ,μ, σ, τ, q_ON_*|***Y***), is related to likelihoods and priors through the expression:

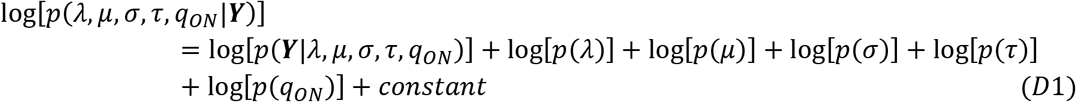

where log[*p*(***Y***|*λ, μ, σ, τ, q_ON_*)] ≡ log*L*(*λ, μ, σ, τ, q_ON_*|***Y***) is the log-likelihood function from Equation (C30) and log[*p*(*λ*)] represents the prior probability function of *λ* (and similarly for the other parameters).

The posterior distribution is estimated using numerical sampling methods, such as Markov chain Monte Carlo (MCMC) algorithms (Stauffer, 2007). They consist in drawing parameter values iteratively, producing a long sequence (chain) of values that converge to the desired distribution. Each newly sampled combination of parameter values will depend on the previous parameter values and on the posterior function (e.g., Equation D1) evaluated at that combination (the higher the function value, the higher the chance that any new combination will be chosen in the sequence). The specifics of how new values are generated depends on the algorithm. Here we used the slice sampling algorithm (Neal, 2003) for its conceptual simplicity and easiness of implementation. It is based on the principle that any probability function can be unbiasedly sampled if we sample uniformly the area under its function (or any function proportional to it, so the *constant* term in Equation D1 can be ignored). The algorithm works by sequentially “slicing” the function firstly with vertical line segments below it, then horizontal segments that determine new parameter values. One advantage of this procedure is that it does not require the use of a proposal distribution for generating parameter values, as in the more commonly used Metropolis-Hastings algorithm (Stauffer, 2007). However, it requires determining before-hand for each parameter the maximum width of the horizontal segment used to search for new values. In principle, the width used for each parameter should be proportional or closely associated to the parameter’s domain in the function (Neal, 2003). To determine the widths, we implemented an adaptive estimation phase based on the automated interval width selection algorithm proposed by Tibbits et al. (2014) as an initial step within the *slicesample* function in MATLAB (Copyright 2005-2011 The MathWorks, Inc).

The prior distributions for all parameters were assumed to be all uniform and non-informative. The prior for *λ* was uniform in the range [-1000,1000]; for the retention selectivity parameters, the ranges were: [−1000,1000] for *μ*, [0,1000] for *σ*, and [0,1] for *τ*. The ON relative catchability coefficient *q_ON_* was estimated in a log_10_-scale, whose prior distribution was uniform in the range [-1000,1000]. Two other parameters determining the size distributions were the minimum (*y_min_* or *x_min_*) and maximum size (*y_max_* or *x_max_*). They were not estimated but imposed as constants in all analyses. For the community data analyses, the size limits (based on length, mm) were group-specific: [*y_min_*, *y_max_*] = [2^−7.875^,2^−1.875^] for phytoplankton, [2^−1.875^, 2^1.125^] for zooplankton, and [2^5.875^, 2^10.375^] for fish (Supplementary Material C). For the gillnet data, the limits were based on ln(weight) (g) and applied to both gears: [*x_min_, x_max_*] = [ln(0.029), ln(94664)] *=* [−3.5398,11.4581]. They were the minimum and maximum observed weights from the entire BsM (Cycles 1 and 2) dataset after excluding outliers (see Supplementary Material A).

All analyses were run in MATLAB R2020b. For each estimation of a joint posterior distribution, we ran 3 independent MCMC chains each with a sample size of 1000 values for each parameter, after a burn-in of 1000 and thinning of 10. The chains were tested for convergence to a stationary distribution by means of visual inspection of traces and through the Gelman-Rubin scale reduction factor R (Gelman and Rubin, 1992), which was lower than 1.1 for all parameters in all analyses. The traces from MCMC runs for all parameters are shown in Figures D1–D13.

**Figure D1.**
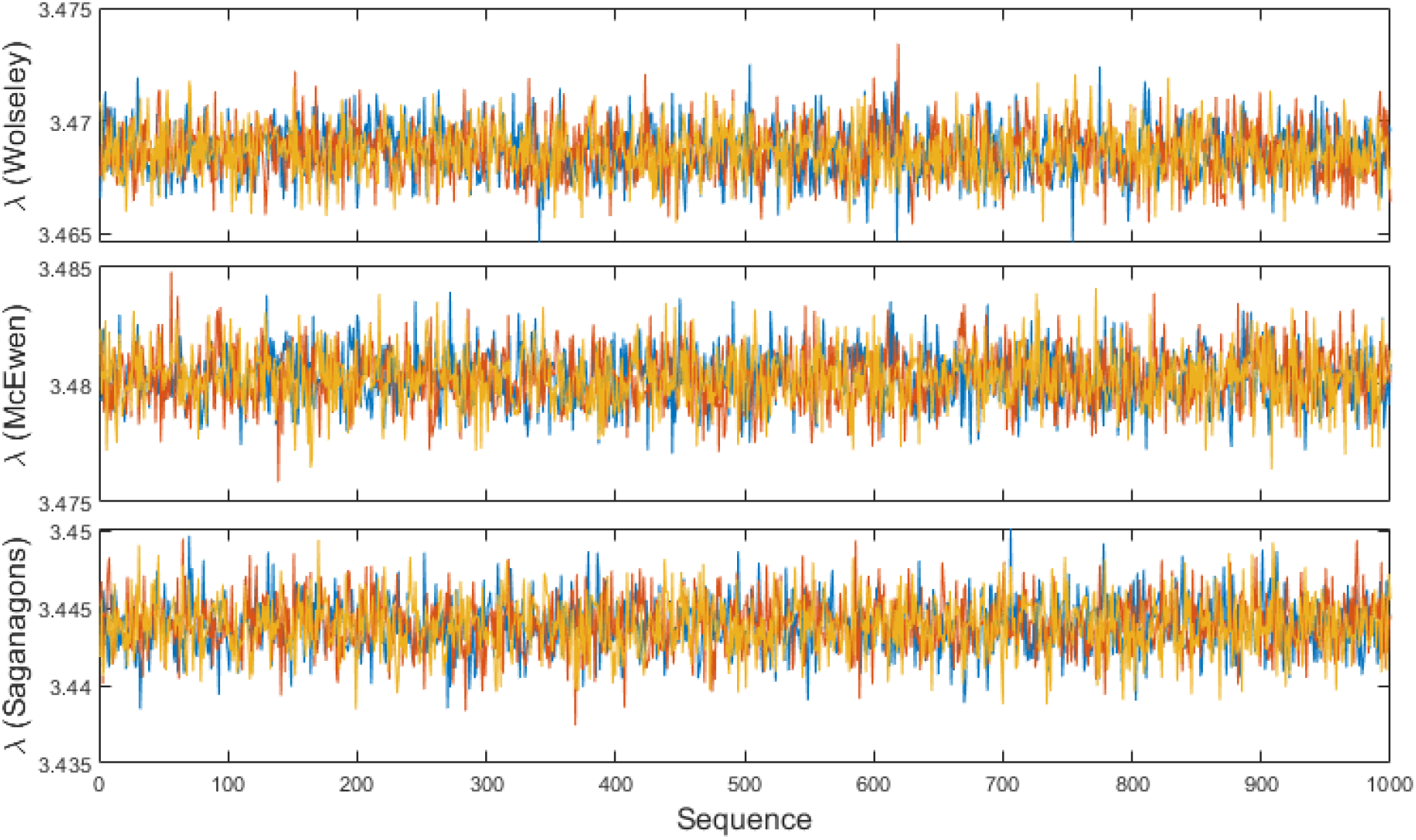
Traces of the size spectrum slope magnitude λ for the PZF community data grouping (phytoplankton + zooplankton + fish). Each color represents one of three independent Bayesian chains.

**Figure D2.**
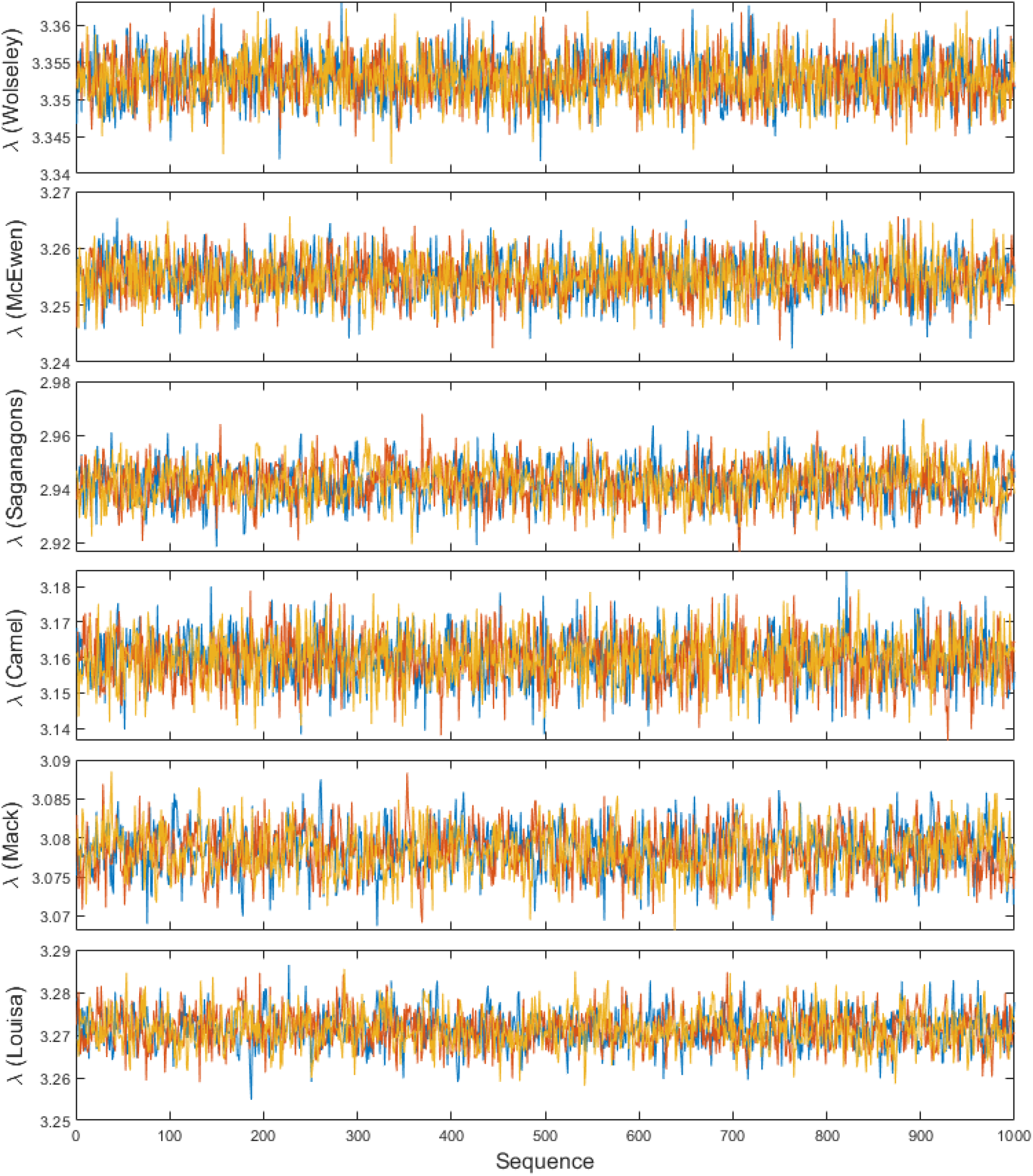
Traces of the size spectrum slope magnitude λ for the ZF community data grouping (zooplankton + fish). Each color represents one of three independent Bayesian chains.

**Figure D3.**
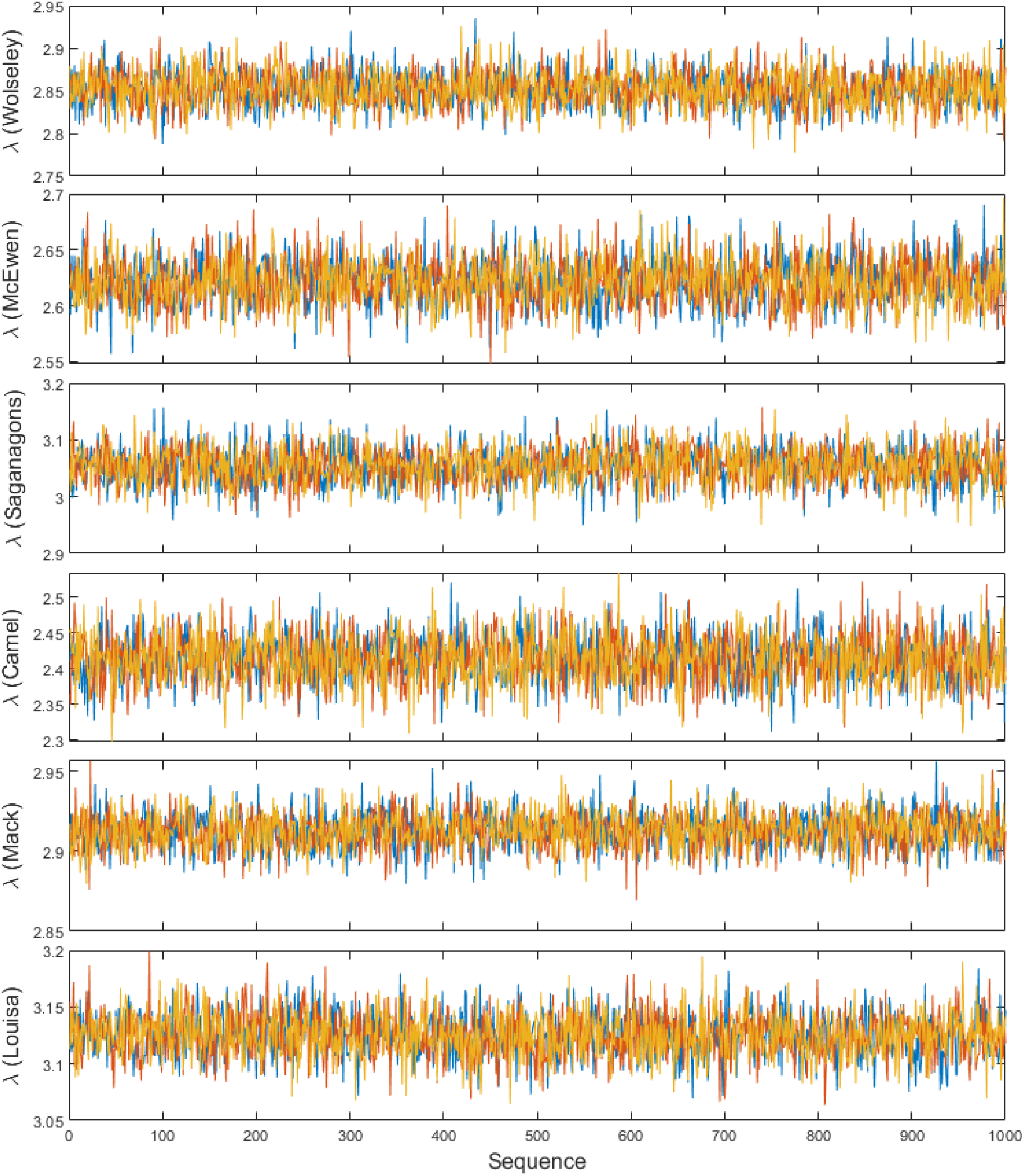
Traces of the size spectrum slope magnitude λ for the F community data grouping (fish). Each color represents one of three independent Bayesian chains.

**Figure D4.**
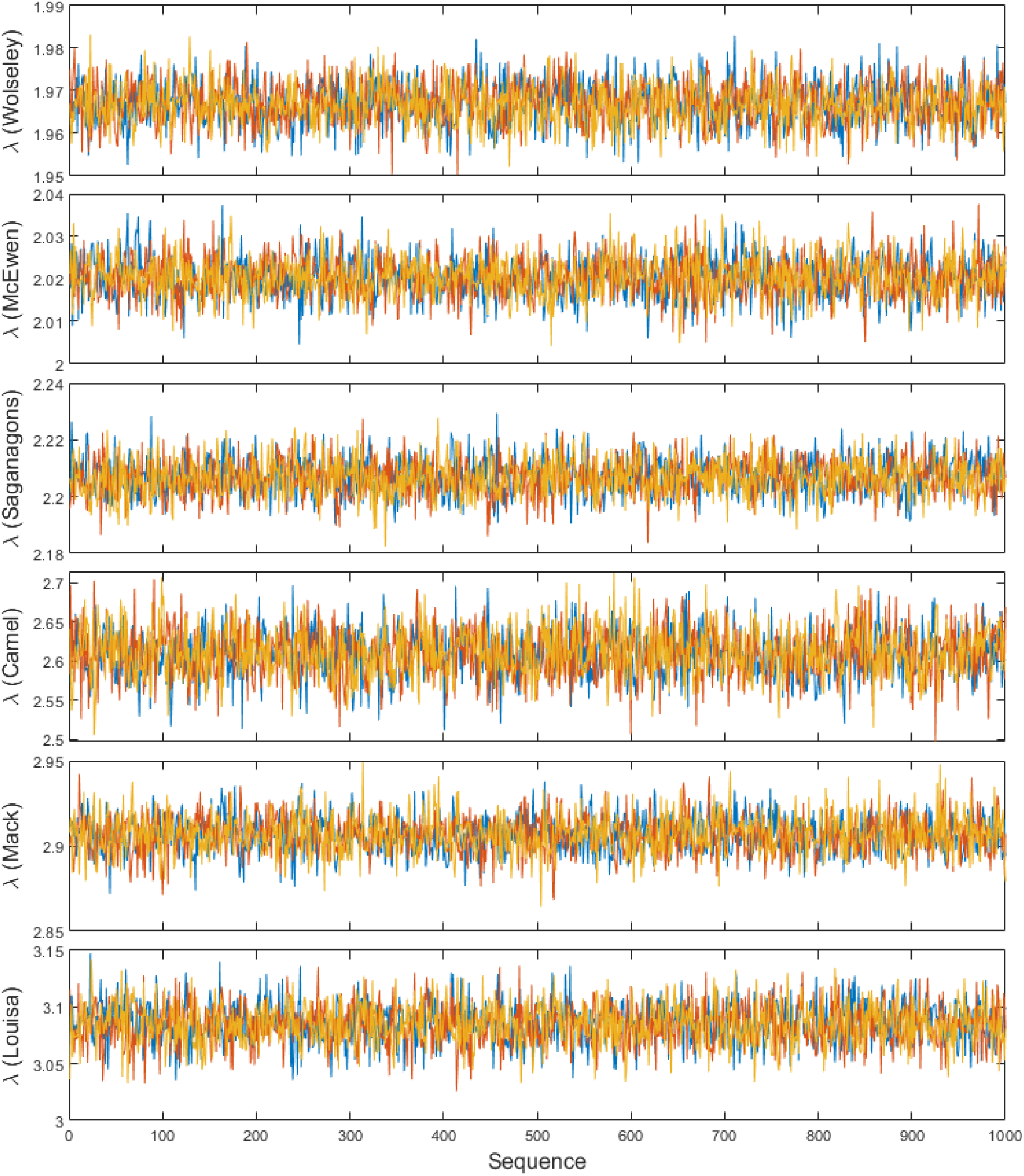
Traces of the size spectrum slope magnitude λ based on within-group variation in the community data. Each color represents one of three independent Bayesian chains.

**Figure D5.**
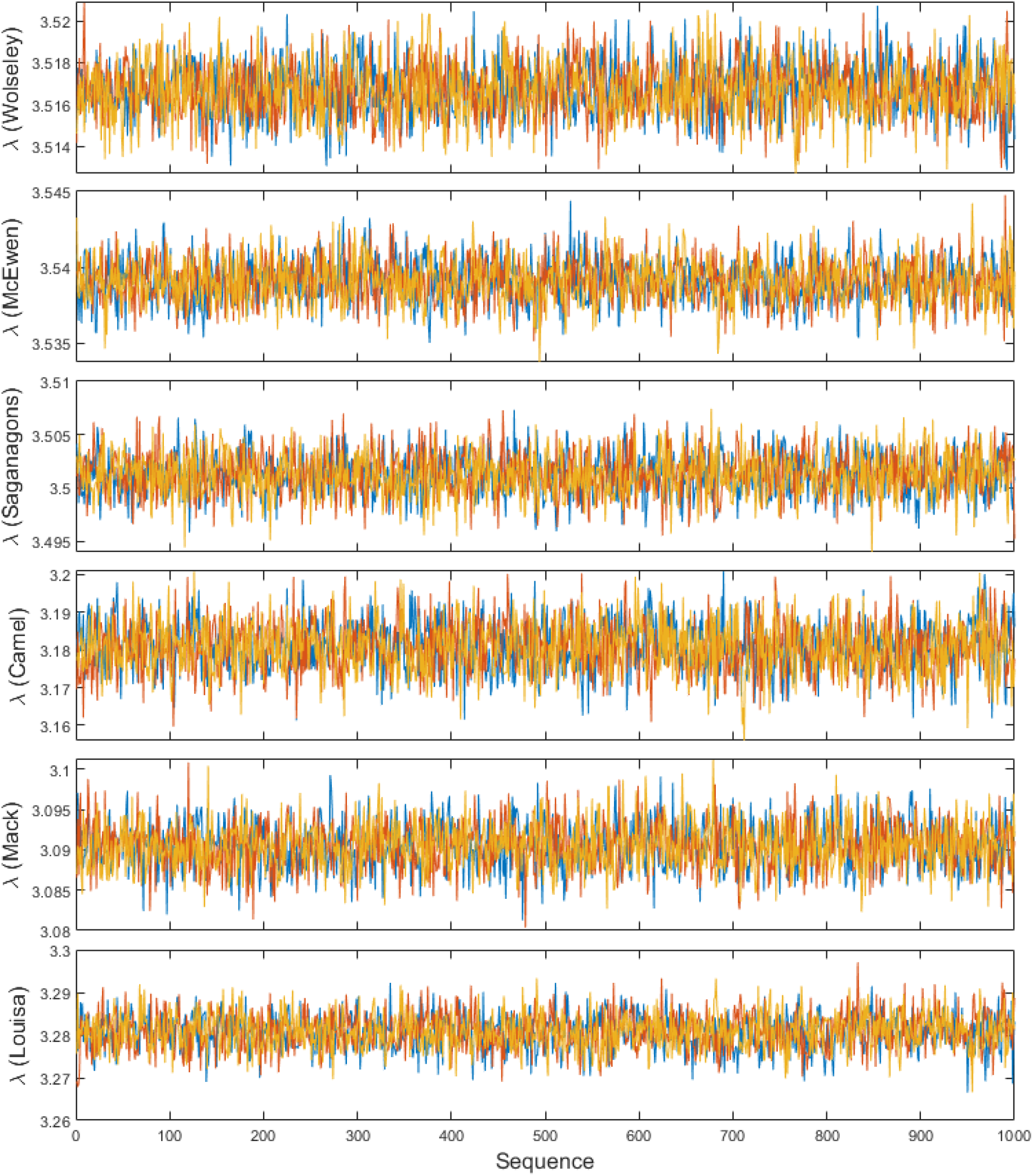
Traces of the size spectrum slope magnitude λ based on between-group variation in the community data. Each color represents one of three independent Bayesian chains.

**Figure D6.**
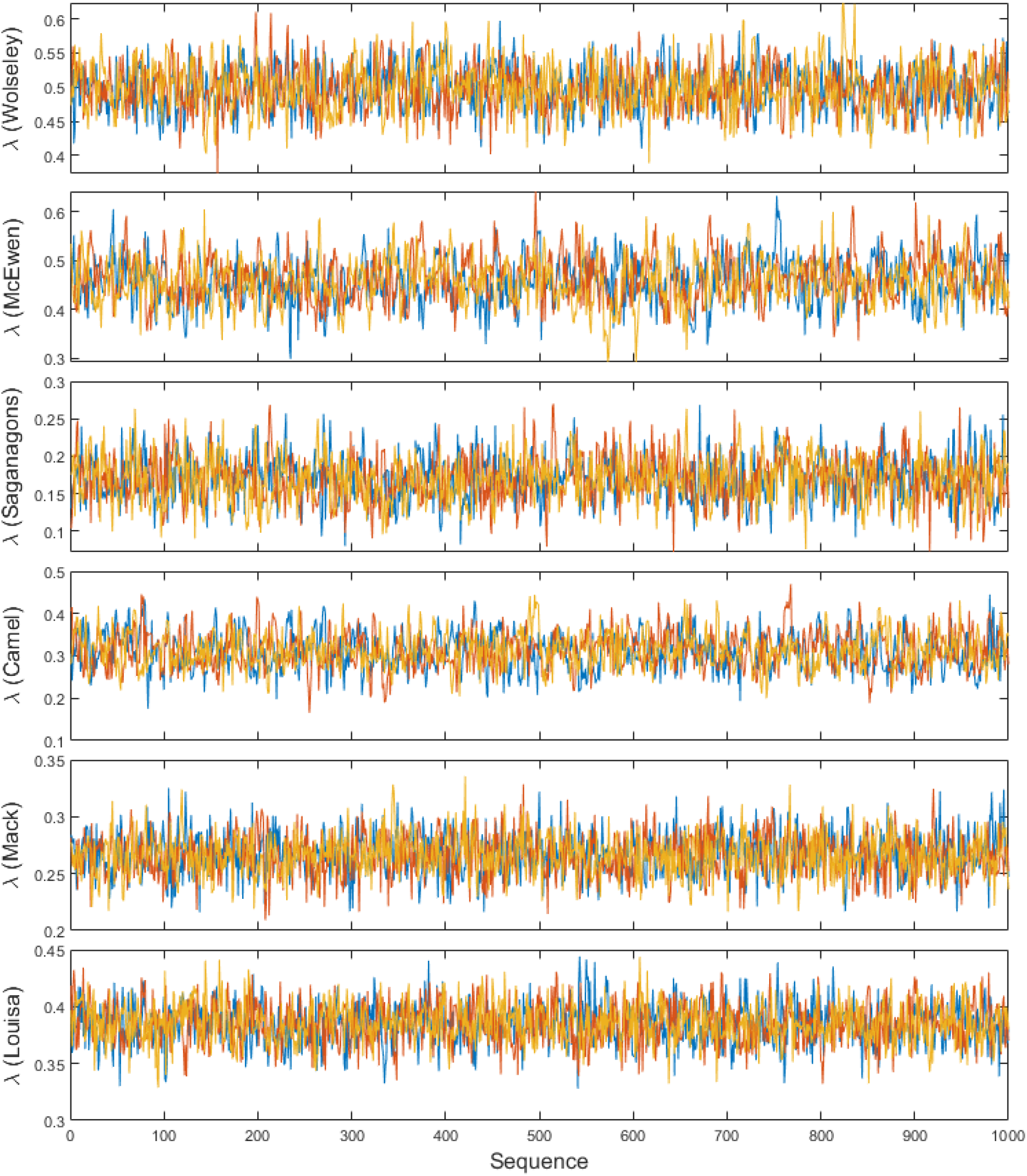
Traces of the size spectrum slope magnitude λ for the ON gillnet data. Each color represents one of three independent Bayesian chains.

**Figure D7.**
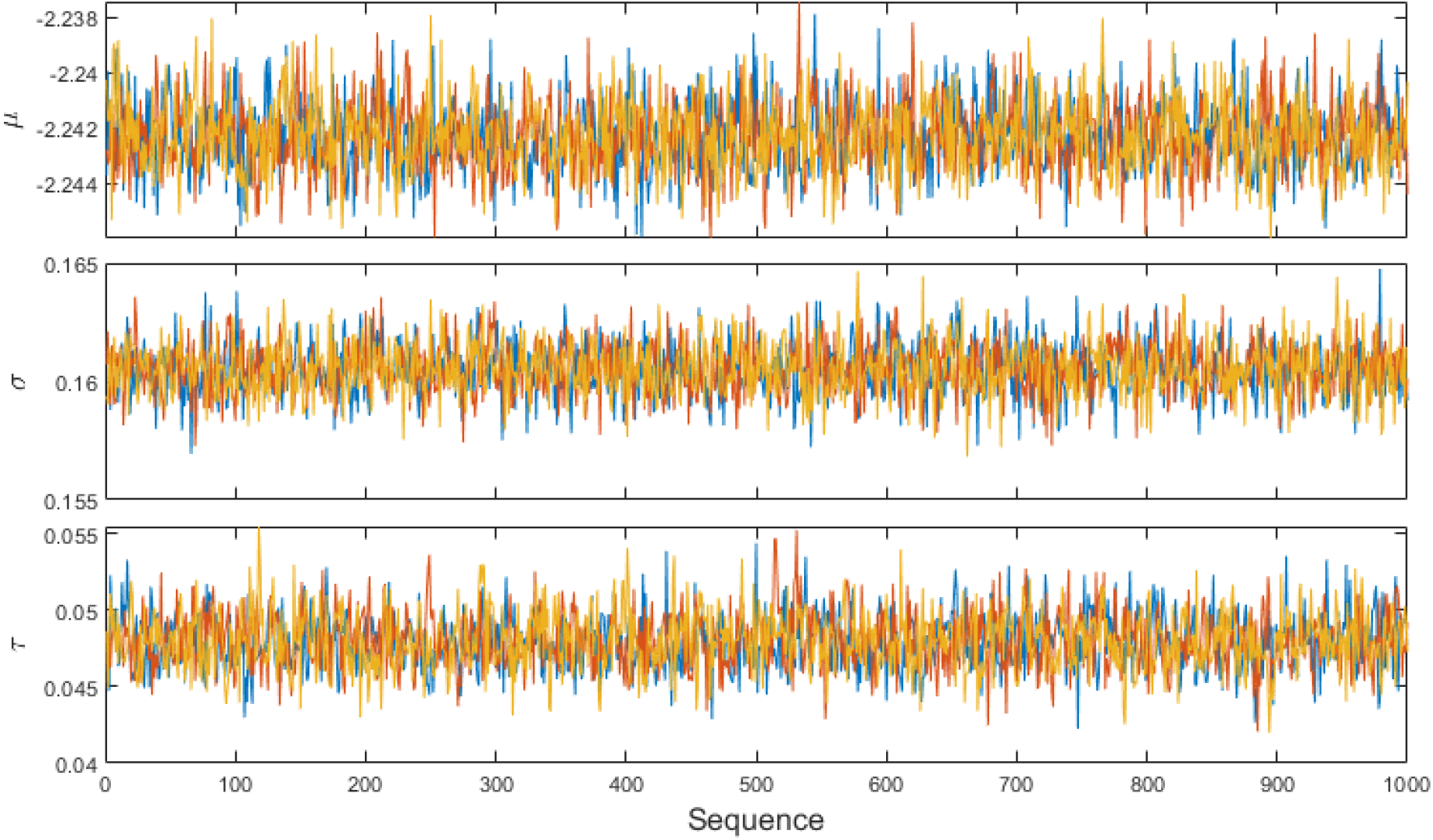
Traces of the retention selectivity parameters, jointly estimated with the ON gillnet data. Each color represents one of three independent Bayesian chains.

**Figure D8.**
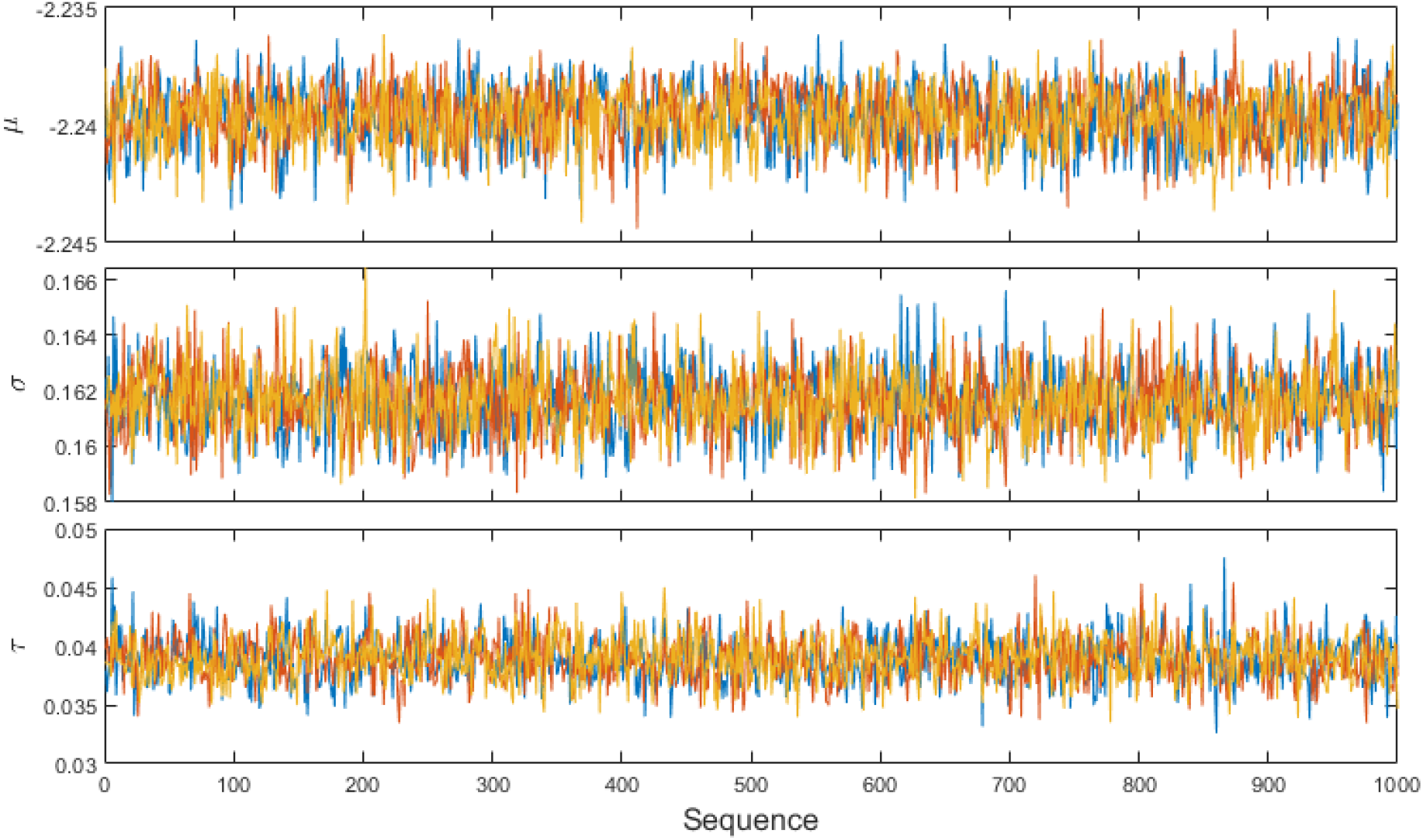
Traces of the retention selectivity parameters, jointly estimated with the NA gillnet data. Each color represents one of three independent Bayesian chains.

**Figure D9.**
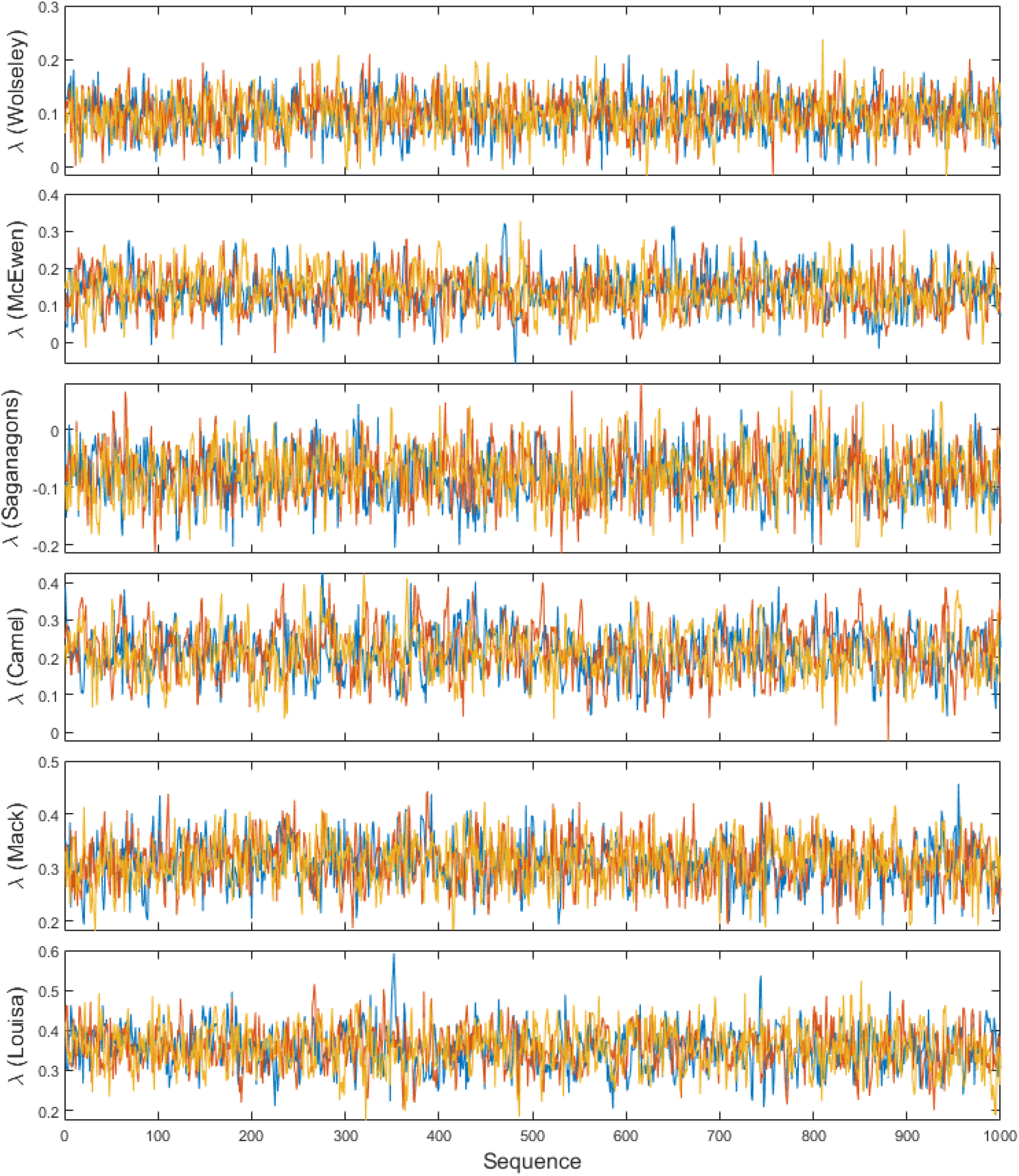
Traces of the size spectrum slope magnitude λ for the NA gillnet data. Each color represents one of three independent Bayesian chains.

**Figure D10.**
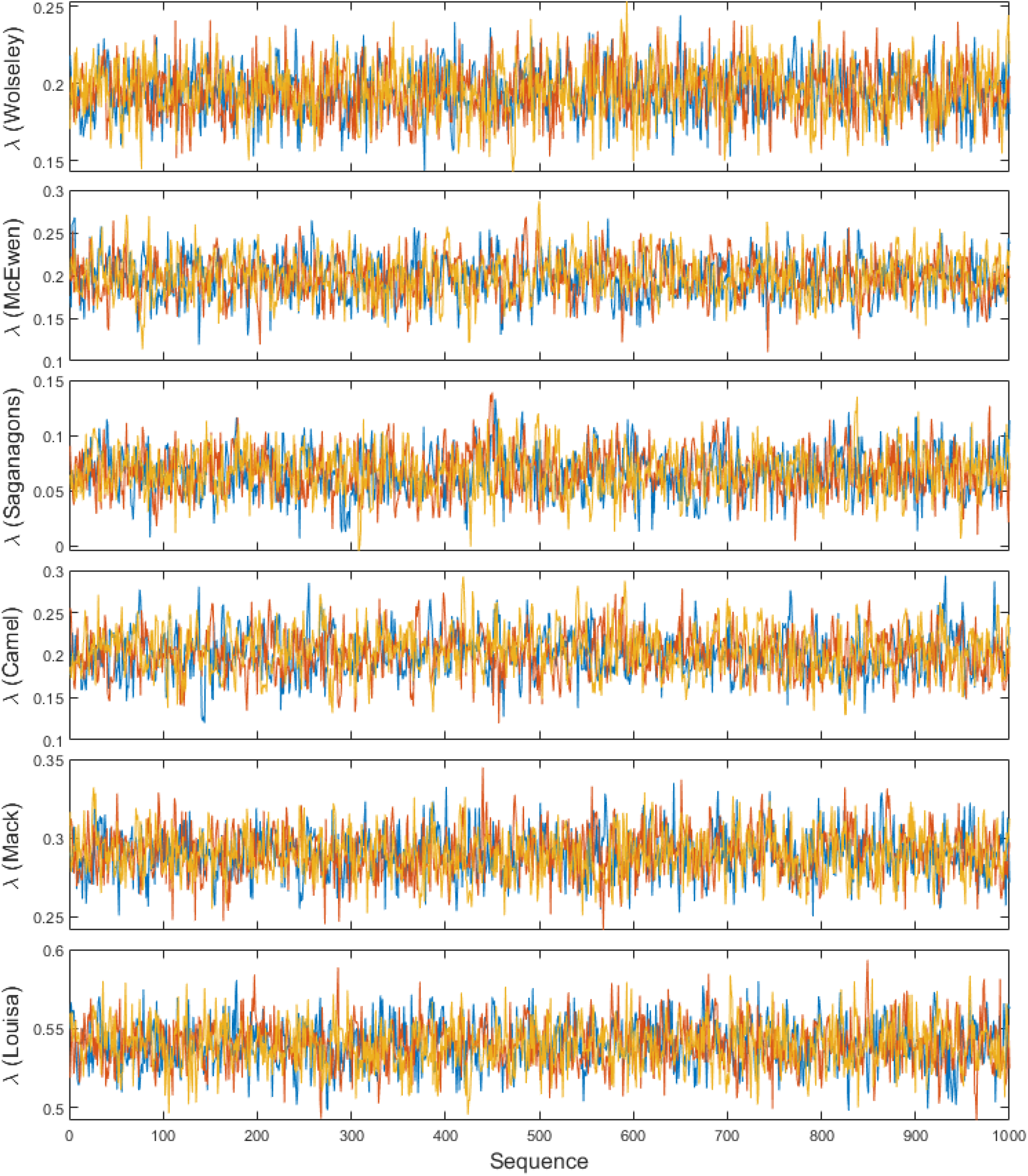
Traces of the size spectrum slope magnitude λ for the BsM (ON + NA) gillnet data. Each color represents one of three independent Bayesian chains.

**Figure D11.**
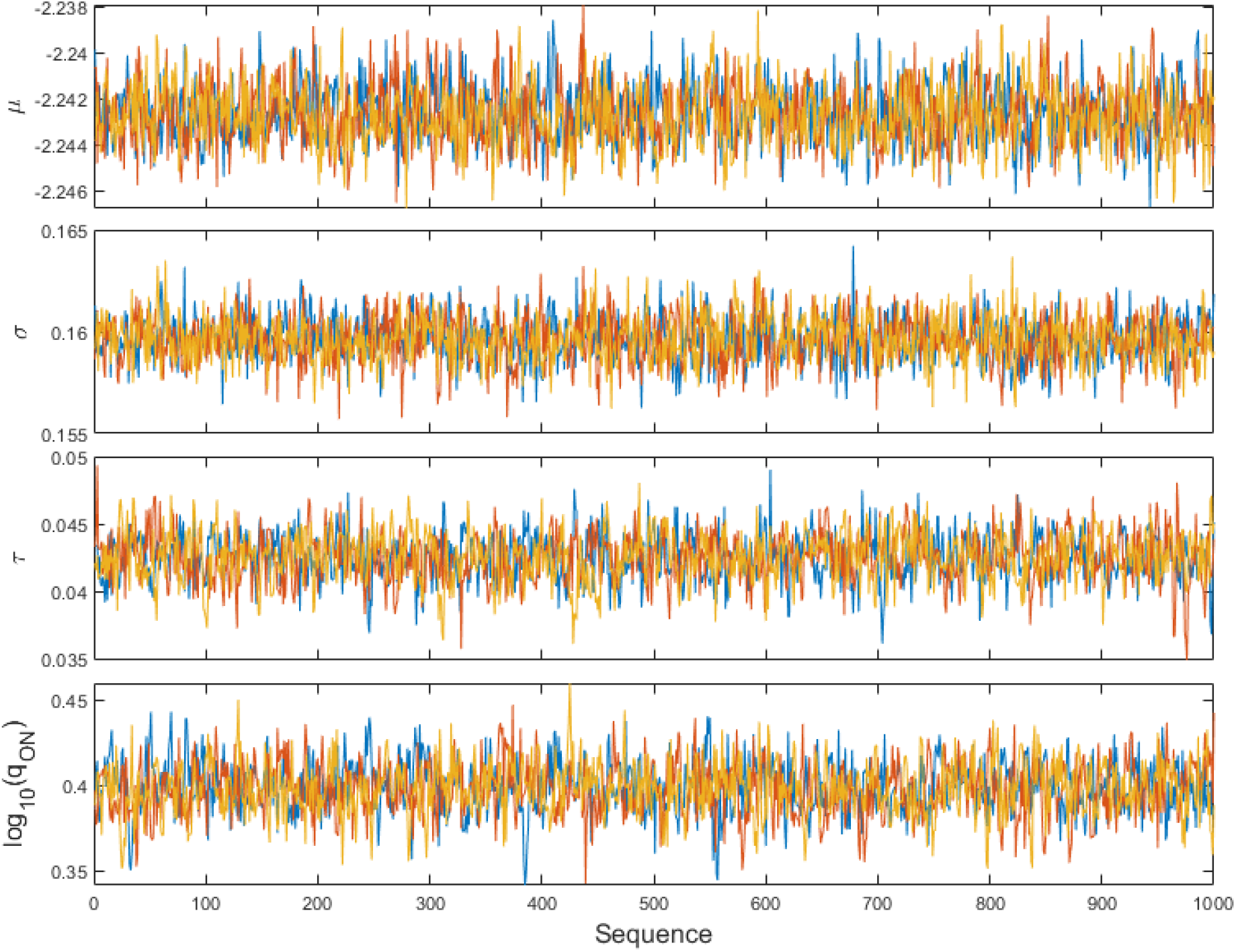
Traces of the retention selectivity parameters and relative ON catchability, jointly estimated with the BsM (ON + NA) gillnet data. Each color represents one of three independent Bayesian chains.

**Figure D12.**
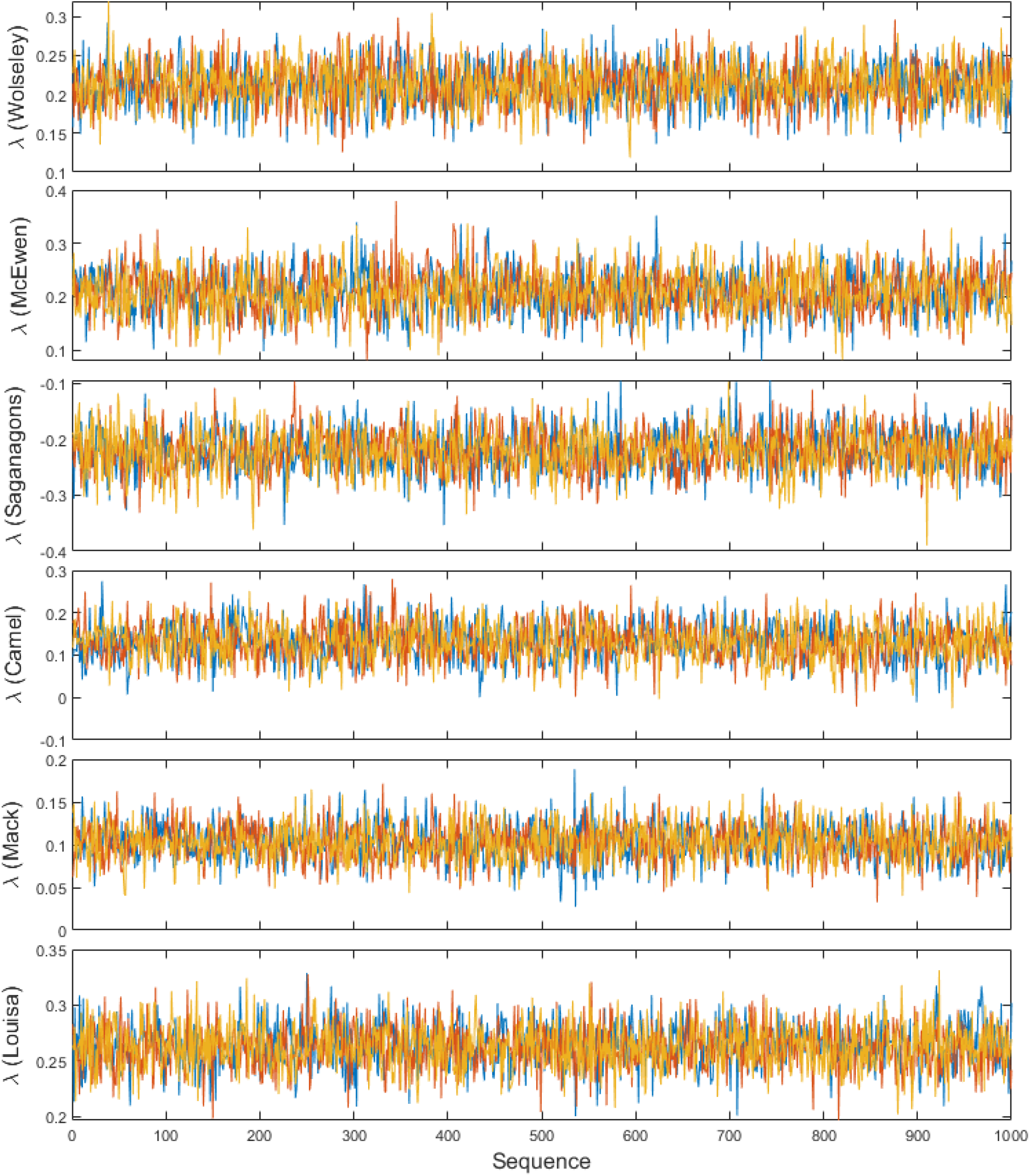
Traces of the size spectrum slope magnitude λ based on within-group variation in the gillnet data. Each color represents one of three independent Bayesian chains.

**Figure D13.**
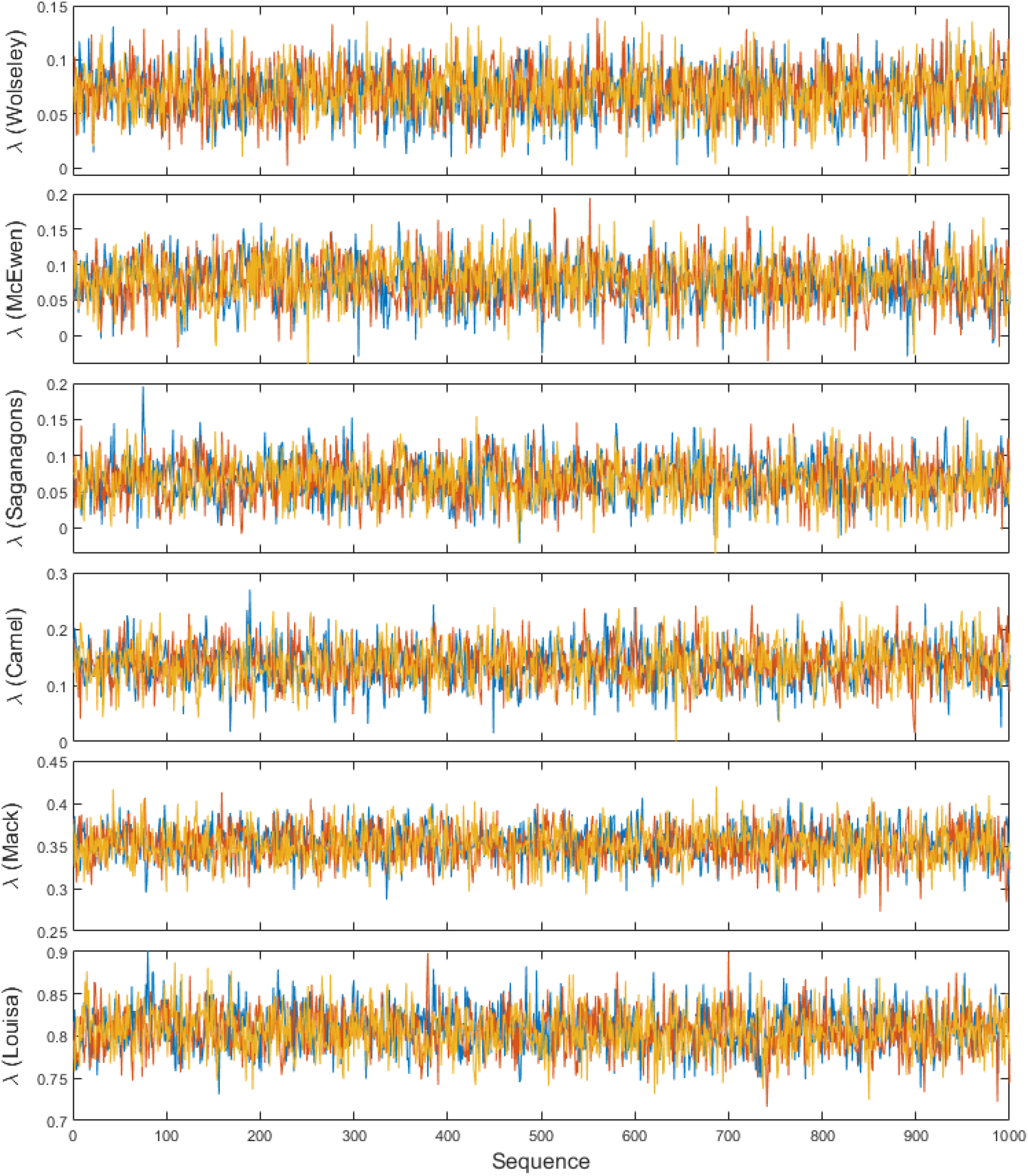
Traces of the size spectrum slope magnitude λ based on between-group variation in the gillnet data. Each color represents one of three independent Bayesian chains.

### Supplementary Material E: additional results

Estimates (percentiles) of all size spectrum parameters and the jointly estimated selectivity parameters are presented in Table E1. The fitted size spectra are plotted with observed counts and expected abundances from the community data in Figure E1 (for all three groups together, phytoplankton + zooplankton + fish, PZF), Figure E2 (zooplankton + fish, ZF), and Figure E3 (fish only, F). In all cases, a strong non-linear secondary structure leads to deviations of observed distributions from the predicted by a power-law size spectrum. For the three lakes with phytoplankton data, the fitted spectrum overestimates the abundances of zooplankton (Figure E1). This could be because of trophic cascades affecting entire groups or because of zooplankton sampling insufficiency that was unaccounted for. The fit seems to be best for ZF communities (Figure E2) and becomes worse when only fish is included (Figure E3), as the secondary structure gets relatively more influential within a shorter size range.

The NA gillnet weight distributions also show important deviations from predictions (Figure E4). One consistent deviation is a concentration of catches around ln(weight) = 7, which is approximately 1kg. This is likely not due to unaccounted gear selectivity as there is no indication that the selectivity model underestimates catches around that weight (Figure B3 right panel, ln(effective length)≈2.3). Two plausible explanations are (i) the low predation mortality experienced by large fish, which bumps up their abundance relative to the power-law expectation (which assumes self-similar predation across the entire size scale) and (ii) a concentration of fish reaching maturity, causing growth to slow down and an accumulation of biomass around that size range. Some of the large species caught in the study lakes, such as Walleye (*Sander vitreus*), Lake trout (*Salvelinus namaycush*) and Northern Pike (*Esox lucius*), have maturation sizes within that overrepresented size range (ln(weight) ≈ 6 to 8, ≈ 0.4 to 3kg).

**Table E1.** Percentiles of parameter estimates from the Bayesian samples. PZF = phytoplankton + zooplankton + fish; ZF = zooplankton + fish, F = fish; ON = Ontario gillnet standard, NA = North American gillnet standard, BsM = Broad-scale Monitoring protocol (combining ON and NA gears); λ = magnitude of the size spectrum slope (the slope itself is −λ); μ, σ, and τ are the parameters of the Symmetric Exponential function for retention selectivity, determining the position of the peak, the spread, and the tangling component, respectively; qON = relative ON catchability coefficient.

| Data type | Grouping | Parameter | Likelihood | Lake | 2.5% | 25% | 50% | 75% | 97.5% |
| --- | --- | --- | --- | --- | --- | --- | --- | --- | --- |
| Community | PZF | $\lambda$ | Full | Wolseley | 3.466 | 3.468 | 3.469 | 3.469 | 3.471 |
| Community | PZF | $\lambda$ | Full | McEwen | 3.478 | 3.479 | 3.480 | 3.481 | 3.483 |
| Community | PZF | $\lambda$ | Full | Saganagons | 3.440 | 3.443 | 3.444 | 3.445 | 3.447 |
| Community | ZF | $\lambda$ | Full | Wolseley | 3.347 | 3.351 | 3.353 | 3.355 | 3.359 |
| Community | ZF | $\lambda$ | Full | McEwen | 3.248 | 3.252 | 3.255 | 3.257 | 3.262 |
| Community | ZF | $\lambda$ | Full | Saganagons | 2.928 | 2.937 | 2.942 | 2.947 | 2.956 |
| Community | ZF | $\lambda$ | Full | Camel | 3.146 | 3.155 | 3.160 | 3.164 | 3.173 |
| Community | ZF | $\lambda$ | Full | Mack | 3.073 | 3.076 | 3.078 | 3.080 | 3.084 |
| Community | ZF | $\lambda$ | Full | Louisa | 3.263 | 3.269 | 3.271 | 3.274 | 3.280 |
| Community | F | $\lambda$ | Full | Wolseley | 2.812 | 2.838 | 2.852 | 2.867 | 2.896 |
| Community | F | $\lambda$ | Full | McEwen | 2.581 | 2.607 | 2.620 | 2.635 | 2.663 |
| Community | F | $\lambda$ | Full | Saganagons | 2.993 | 3.032 | 3.054 | 3.076 | 3.116 |
| Community | F | $\lambda$ | Full | Camel | 2.349 | 2.391 | 2.413 | 2.436 | 2.478 |
| Community | F | $\lambda$ | Full | Mack | 2.891 | 2.905 | 2.913 | 2.920 | 2.935 |
| Community | F | $\lambda$ | Full | Louisa | 3.087 | 3.112 | 3.124 | 3.138 | 3.164 |
| Community | PZF | $\lambda$ | Within-group | Wolseley | 1.957 | 1.964 | 1.967 | 1.970 | 1.976 |
| Community | PZF | $\lambda$ | Within-group | McEwen | 2.011 | 2.017 | 2.020 | 2.023 | 2.030 |
| Community | PZF | $\lambda$ | Within-group | Saganagons | 2.194 | 2.202 | 2.207 | 2.211 | 2.220 |
| Community | ZF | $\lambda$ | Within-group | Camel | 2.549 | 2.588 | 2.609 | 2.630 | 2.670 |
| Community | ZF | $\lambda$ | Within-group | Mack | 2.885 | 2.899 | 2.907 | 2.914 | 2.929 |
| Community | ZF | $\lambda$ | Within-group | Louisa | 3.048 | 3.073 | 3.085 | 3.097 | 3.121 |
| Community | PZF | $\lambda$ | Between-group | Wolseley | 3.514 | 3.516 | 3.517 | 3.518 | 3.519 |
| Community | PZF | $\lambda$ | Between-group | McEwen | 3.536 | 3.538 | 3.539 | 3.540 | 3.542 |
| Community | PZF | $\lambda$ | Between-group | Saganagons | 3.497 | 3.500 | 3.501 | 3.503 | 3.505 |
| Community | ZF | $\lambda$ | Between-group | Camel | 3.168 | 3.176 | 3.181 | 3.185 | 3.194 |
| Community | ZF | $\lambda$ | Between-group | Mack | 3.085 | 3.089 | 3.091 | 3.092 | 3.096 |
| Community | ZF | $\lambda$ | Between-group | Louisa | 3.273 | 3.278 | 3.281 | 3.284 | 3.289 |
| Gillnet | ON | $\mu$ | Full | | -2.245 | -2.243 | -2.242 | -2.242 | -2.240 |
| Gillnet | ON | $\sigma$ | Full | | 0.158 | 0.160 | 0.161 | 0.161 | 0.163 |
| Gillnet | ON | $\tau$ | Full | | 0.045 | 0.047 | 0.048 | 0.049 | 0.052 |
| Gillnet | ON | $\lambda$ | Full | Wolseley | 0.435 | 0.476 | 0.498 | 0.521 | 0.564 |
| Gillnet | ON | $\lambda$ | Full | McEwen | 0.370 | 0.428 | 0.459 | 0.490 | 0.554 |
| Gillnet | ON | $\lambda$ | Full | Saganagons | 0.111 | 0.149 | 0.170 | 0.189 | 0.233 |
| Gillnet | ON | $\lambda$ | Full | Camel | 0.234 | 0.284 | 0.312 | 0.341 | 0.403 |
| Gillnet | ON | $\lambda$ | Full | Mack | 0.230 | 0.254 | 0.267 | 0.279 | 0.302 |
| Gillnet | ON | $\lambda$ | Full | Louisa | 0.350 | 0.373 | 0.386 | 0.397 | 0.420 |
| Gillnet | NA | $\mu$ | Full | | -2.242 | -2.241 | -2.240 | -2.239 | -2.237 |
| Gillnet | NA | $\sigma$ | Full | | 0.159 | 0.161 | 0.162 | 0.162 | 0.164 |
| Gillnet | NA | $\tau$ | Full | | 0.035 | 0.038 | 0.039 | 0.040 | 0.043 |
| Gillnet | NA | $\lambda$ | Full | Wolseley | 0.024 | 0.073 | 0.098 | 0.123 | 0.168 |
| Gillnet | NA | $\lambda$ | Full | McEwen | 0.035 | 0.101 | 0.137 | 0.172 | 0.247 |
| Gillnet | NA | $\lambda$ | Full | Saganagons | -0.163 | -0.108 | -0.079 | -0.048 | 0.009 |
| Gillnet | NA | $\lambda$ | Full | Camel | 0.091 | 0.168 | 0.210 | 0.256 | 0.337 |
| Gillnet | NA | $\lambda$ | Full | Mack | 0.224 | 0.280 | 0.310 | 0.338 | 0.394 |
| Gillnet | NA | $\lambda$ | Full | Louisa | 0.254 | 0.319 | 0.355 | 0.392 | 0.455 |
| Gillnet | BsM | $\mu$ | Full | | -2.245 | -2.244 | -2.243 | -2.242 | -2.240 |
| Gillnet | BsM | $\sigma$ | Full | | 0.157 | 0.159 | 0.160 | 0.160 | 0.162 |
| Gillnet | BsM | $\tau$ | Full | | 0.039 | 0.041 | 0.043 | 0.044 | 0.046 |
| Gillnet | BsM | $q_{ON}$ | Full | | 2.330 | 2.445 | 2.504 | 2.563 | 2.685 |
| Gillnet | BsM | $\lambda$ | Full | Wolseley | 0.162 | 0.184 | 0.195 | 0.206 | 0.227 |
| Gillnet | BsM | $\lambda$ | Full | McEwen | 0.151 | 0.181 | 0.197 | 0.213 | 0.245 |
| Gillnet | BsM | $\lambda$ | Full | Saganagons | 0.029 | 0.053 | 0.066 | 0.080 | 0.106 |
| Gillnet | BsM | $\lambda$ | Full | Camel | 0.154 | 0.186 | 0.203 | 0.220 | 0.256 |
| Gillnet | BsM | $\lambda$ | Full | Mack | 0.262 | 0.280 | 0.289 | 0.300 | 0.319 |
| Gillnet | BsM | $\lambda$ | Full | Louisa | 0.511 | 0.530 | 0.539 | 0.550 | 0.569 |
| Gillnet | BsM | $\lambda$ | Within-group | Wolseley | 0.157 | 0.193 | 0.212 | 0.230 | 0.265 |
| Gillnet | BsM | $\lambda$ | Within-group | McEwen | 0.131 | 0.180 | 0.208 | 0.236 | 0.287 |
| Gillnet | BsM | $\lambda$ | Within-group | Saganagons | -0.294 | -0.246 | -0.221 | -0.196 | -0.147 |
| Gillnet | BsM | $\lambda$ | Within-group | Camel | 0.046 | 0.102 | 0.131 | 0.160 | 0.216 |
| Gillnet | BsM | $\lambda$ | Within-group | Mack | 0.061 | 0.088 | 0.103 | 0.117 | 0.145 |
| Gillnet | BsM | $\lambda$ | Within-group | Louisa | 0.224 | 0.249 | 0.262 | 0.276 | 0.301 |
| Gillnet | BsM | $\lambda$ | Between-group | Wolseley | 0.026 | 0.055 | 0.070 | 0.086 | 0.115 |
| Gillnet | BsM | $\lambda$ | Between-group | McEwen | 0.011 | 0.055 | 0.077 | 0.099 | 0.138 |
| Gillnet | BsM | $\lambda$ | Between-group | Saganagons | 0.011 | 0.047 | 0.066 | 0.085 | 0.124 |
| Gillnet | BsM | $\lambda$ | Between-group | Camel | 0.070 | 0.114 | 0.137 | 0.161 | 0.208 |
| Gillnet | BsM | $\lambda$ | Between-group | Mack | 0.311 | 0.338 | 0.351 | 0.366 | 0.391 |
| Gillnet | BsM | $\lambda$ | Between-group | Louisa | 0.758 | 0.789 | 0.807 | 0.824 | 0.858 |

**Figure E1.**
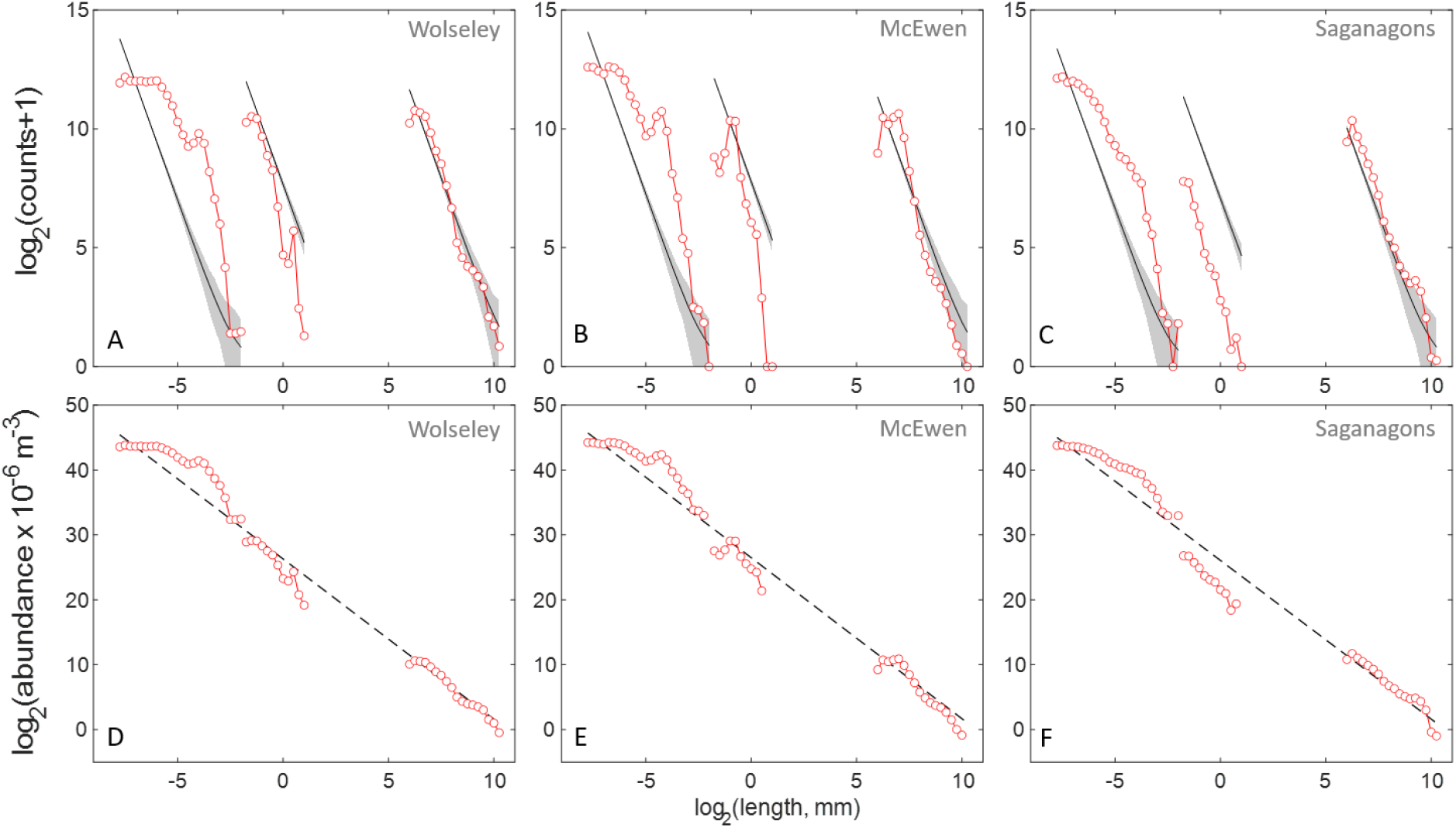
Observed (red lines and dots) and predicted (gray bands = 95% prediction intervals, black lines = median) (A-C) individual counts from community data samples and (D-F) expected abundances (individuals per million m^3^) of phytoplankton, zooplankton, and fish (PZF) from the three lakes with complete data.

**Figure E2.**
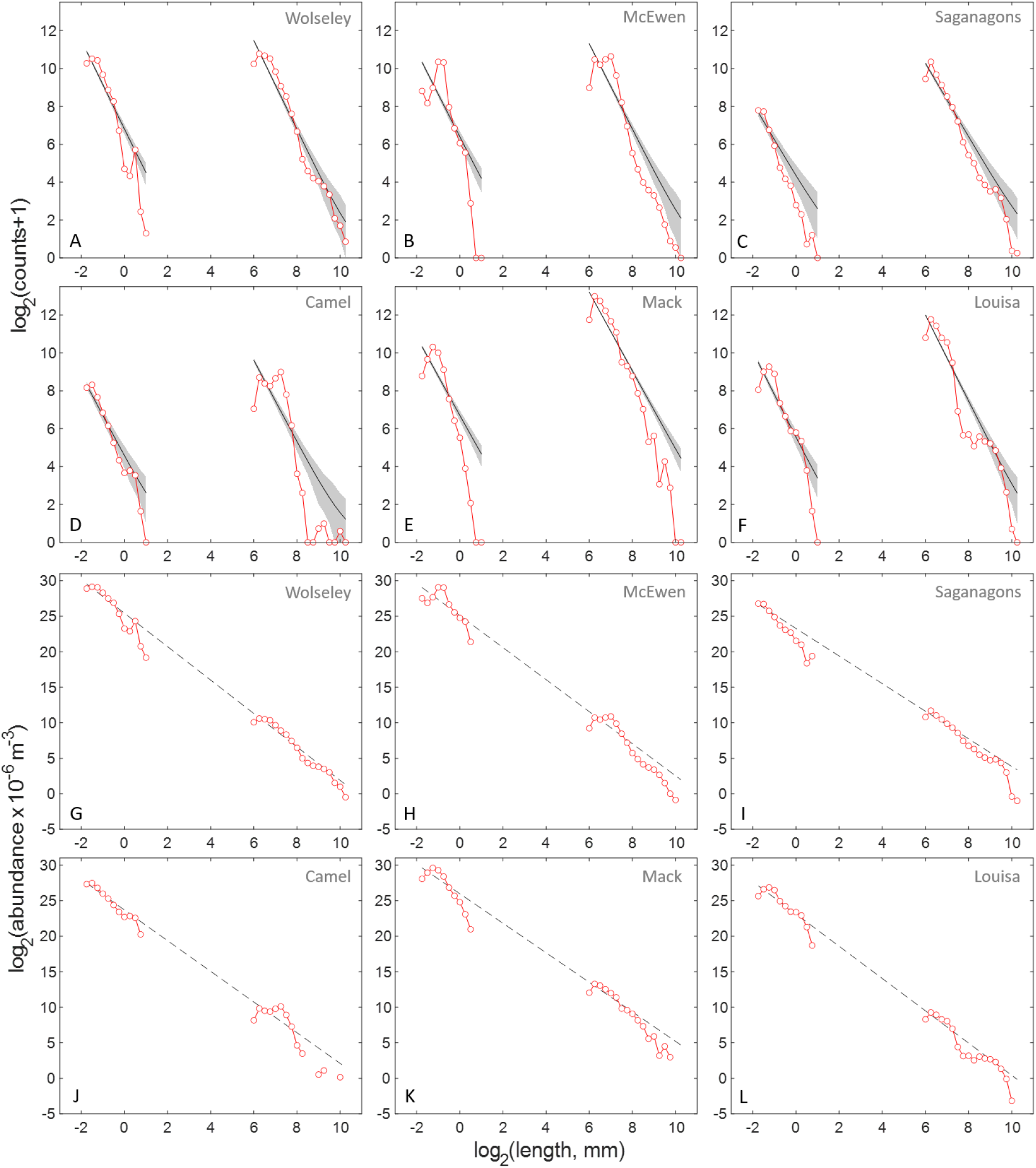
Observed (red lines and dots) and predicted (gray bands = 95% prediction intervals, black lines = median) (A-F) individual counts from community data samples and (G-L) expected abundances (individuals per million m^3^) of zooplankton and fish (ZF) from the six lakes.

**Figure E3.**
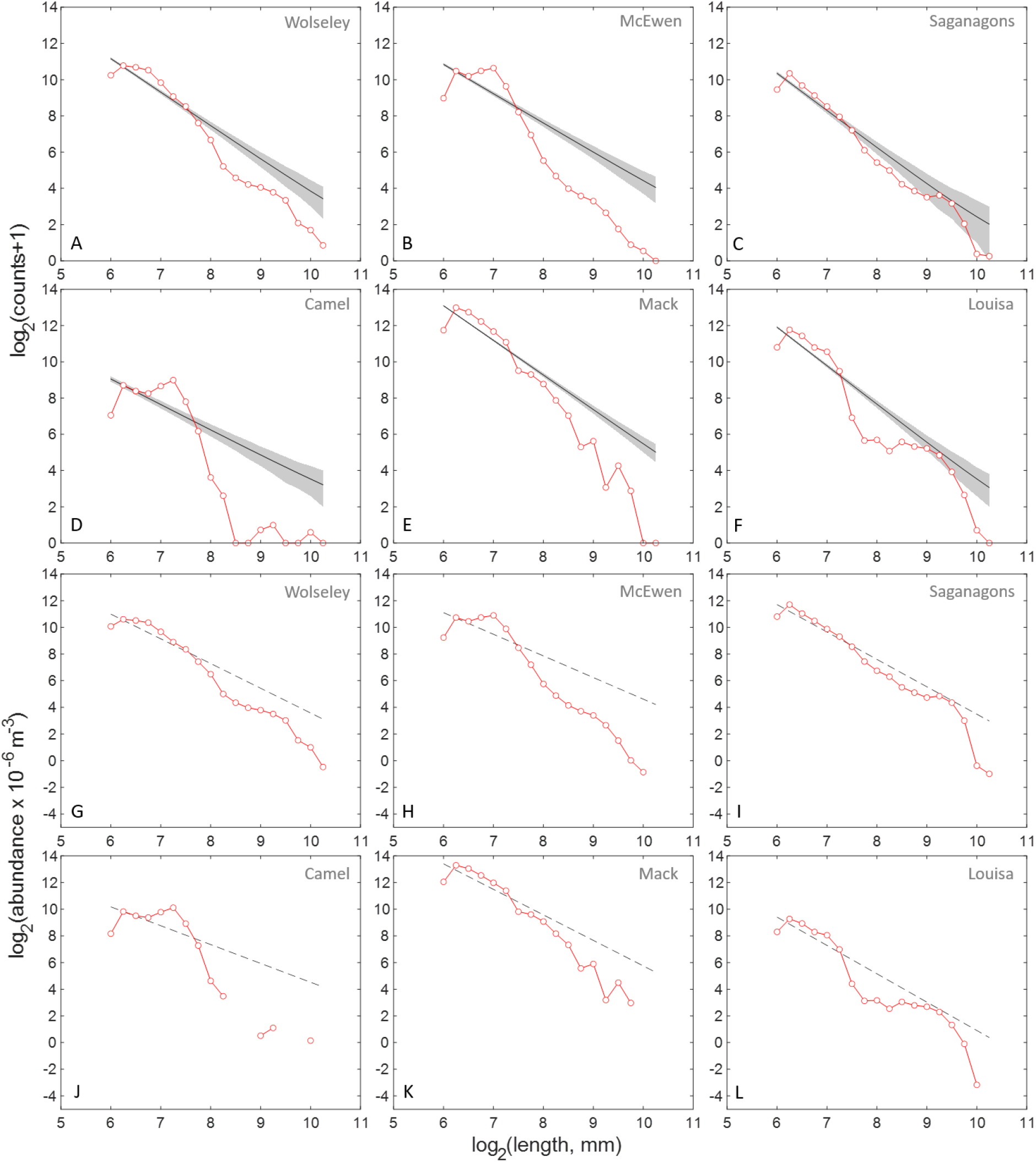
Observed (red lines and dots) and predicted (gray bands = 95% prediction intervals, black lines = median) (A-F) individual counts from community data samples and (G-L) expected abundances (individuals per million m^3^) of fish (F) from the six lakes.

**Figure E4.**
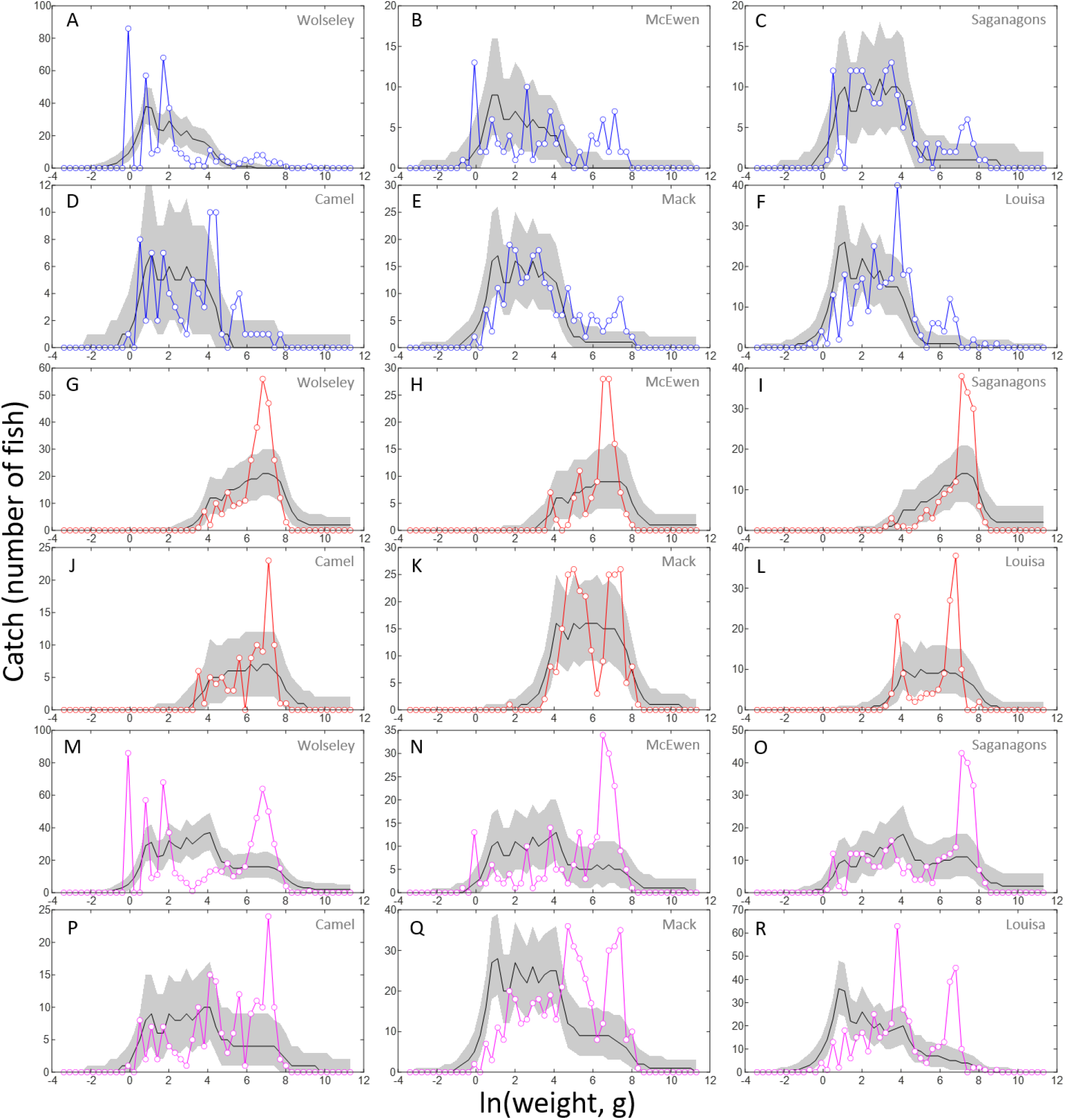
Observed (colored lines and dots) and predicted (gray bands = 95% prediction intervals, black lines = median) catch from gillnet surveys. A-F (blue lines): ON gillnets. G-L (red lines): NA gillnets. M-R (purple lines): BsM (all nets combined). Body size (*x* = ln (*W*)) was divided into 50 equally spaced bins. Predictions were generated from 10 Multinomial distributed replicate values for each one of the 3000 Bayesian samples. The multinomial probability parameters were calculated by integrating the probability function *o_g_*(*x*) (Equation C20) for individual gear distributions (A-L) or *o*(*x*) (Equation C25) for the combined BsM distributions (M-R) over each bin’s range and multiplying by the total catch from the respective gillnet gear combination in each lake.

## References

Andersen, K.H., Beyer, J.E., 2006. Asymptotic size determines species abundance in the marine size spectrum. The American Naturalist, 168(1), 54–61.

Andersen, K.H., Jacobsen, N.S., Farnsworth, K.D., 2016. The theoretical foundations for size spectrum models of fish communities. Can. J. Fish. Aquat. Sci., 73(4), 575–588.

Anderson, C.S., 1998. Partitioning total size selectivity of gill nets for walleye (Stizostedion vitreum) into encounter, contact, and retention components. Can. J. Fish. Aquat. Sci., 55(8), 1854–1863.

Atkinson, A., Lilley, M.K., Hirst, A.G., McEvoy, A.J., Tarran, G.A., Widdicombe, C., Fileman, E.S., Woodward, E.M.S., Schmidt, K., Smyth, T.J., Somerfield, P.J., 2021. Increasing nutrient stress reduces the efficiency of energy transfer through planktonic size spectra. Limnology and Oceanography, 66, 422–437.

Blanchard, J.L., Jennings, S., Law, R., Castle, M.D., McCloghrie, P., Rochet, M.J., Benoît, E., 2009. How does abundance scale with body size in coupled size-structured food webs? Journal of Animal Ecology, 78(1), 270–280.

Boudreau, P.R., Dickie, L.M., Kerr, S.R., 1991. Body-size spectra of production and biomass as systeml-evel indicators of ecological dynamics. Journal of Theoretical Biology, 152(3), 329–339.

Boudreau, P.R., Dickie, L.M., 1992. Biomass spectra of aquatic ecosystems in relation to fisheries yield. Can. J. Fish. Aquat. Sci., 49(8), 1528–1538.

Braun, L.M., Brucet, S., Mehner, T., 2021. Top-down and bottom-up effects on zooplankton size distribution in a deep stratified lake. Aquatic Ecology, https://doi.org/10.1007/s10452-021-09843-8.

Chang, C.W., Miki, T., Shiah, F.K., Kao, S.J., Wu, J.T., Sastri, A.R., Hsieh, C.H., 2014. Linking secondary structure of individual size distribution with nonlinear size–trophic level relationship in food webs. Ecology, 95(4), 897–909.

Cohen, J.E., Jonsson, T., Carpenter, S.R., 2003. Ecological community description using the food web, species abundance, and body size. Proc. Natl. Acad. Sci. U.S.A., 100(4), 1781–1786.

Datta, S., Blanchard, J.L., 2016. The effects of seasonal processes on size spectrum dynamics. Can. J. Fish. Aquat. Sci., 73(4), 598–610.

Edwards, A.M., Robinson, J.P., Blanchard, J.L., Baum, J.K., Plank, M.J., 2020. Accounting for the bin structure of data removes bias when fitting size spectra. Mar. Ecol. Prog. Ser., 636, 19–33.

Edwards, A.M., Robinson, J.P., Plank, M.J., Baum, J.K., Blanchard, J.L., 2017. Testing and recommending methods for fitting size spectra to data. Methods in Ecology and Evolution, 8(1), 57–67.

Elton, C.S., 2001. Animal Ecology. University of Chicago Press.

Emmrich, M., Brucet, S., Ritterbusch, D., Mehner, T., 2011. Size spectra of lake fish assemblages: responses along gradients of general environmental factors and intensity of lake-use. Freshwater Biology, 56(11), 2316–2333.

Heather, F.J., Blanchard, J.L., Edgar, G.J., Trebilco, R., Stuart-Smith, R.D., 2021. Globally consistent reef size spectra integrating fishes and invertebrates. Ecology Letters, 24(3), 572–579.

Kerr, S.R., Dickie, L.M., 2001. The biomass spectrum: a predator-prey theory of aquatic production. Columbia University Press.

Law, R., Plank, M.J., James, A., Blanchard, J.L., 2009. Size-spectra dynamics from stochastic predation and growth of individuals. Ecology, 90(3), 802–811.

Lester, N.P., Sandstrom, S., de Kerckhove, D.T., Armstrong, K., Ball, H., Amos, J., Dunkley, T., Rawson, M., Addison, P., Dextrase, A. and Taillon, D., Wasylenko, B., Lennox III, P., Giacomini, H.C., Chu, C. 2021. Standardized Broad-Scale Management and Monitoring of Inland Lake Recreational Fisheries: An Overview of the Ontario Experience. Fisheries, 46(3), 107–118.

Mehner, T., Lischke, B., Scharnweber, K., Attermeyer, K., Brothers, S., Gaedke, U., Hilt, S., Brucet, S., 2018. Empirical correspondence between trophic transfer efficiency in freshwater food webs and the slope of their size spectra. Ecology, 99(6), 1463–1472.

Millar, R.B., Fryer, R.J., 1999. Estimating the size-selection curves of towed gears, traps, nets and hooks. Reviews in Fish Biology and Fisheries, 9(1), 89–116.

Neal, R.M., 2003. Slice sampling. Annals of statistics, 705–741.

Ontario Parks. 2019. Quetico Provincial Park Fisheries Management and Aquatic Ecosystem Stewardship Plan: Draft Plan for Discussion. Quetico Provincial Park. 50 p + app.

Perkins, D.M., Durance, I., Jackson, M., Jones, J.I., Lauridsen, R.B., Layer-Dobra, K., Reiss, J., Thompson, M.S., Woodward, G., 2021. Systematic variation in food web body-size structure linked to external subsidies. Biology Letters, 17(3), p.20200798.

Rice, J., Gislason, H., 1996. Patterns of change in the size spectra of numbers and diversity of the North Sea fish assemblage, as reflected in surveys and models. ICES J. Mar. Sci., 53(6), 1214–1225.

Robinson, J.P., Williams, I.D., Edwards, A.M., McPherson, J., Yeager, L., Vigliola, L., Brainard, R.E., Baum, J.K., 2017. Fishing degrades size structure of coral reef fish communities. Global Change Biology, 23(3), 1009–1022.

Rochet, M.J., Benoît, E., 2012. Fishing destabilizes the biomass flow in the marine size spectrum. Proc. Roy. Soc. B: Biological Sciences, 279(1727), 284–292.

Rossberg, A.G., Gaedke, U., Kratina, P., 2019. Dome patterns in pelagic size spectra reveal strong trophic cascades. Nature Communications, 10(1), 1–11.

Rudstam, L.G., Magnuson, J.J., Tonn, W.M., 1984. Size selectivity of passive fishing gear: a correction for encounter probability applied to gill nets. Can. J. Fish. Aquat. Sci., 41(8), 1252–1255.

Sprules, W.G., Barth, L.E., 2016. Surfing the biomass size spectrum: some remarks on history, theory, and application. Can. J. Fish. Aquat. Sci., 73(4), 477–495.

Sprules, W.G., Munawar, M., 1986. Plankton size spectra in relation to ecosystem productivity, size, and perturbation. Can. J. Fish. Aquat. Sci., 43(9), 1789–1794.

Trebilco, R., Baum, J.K., Salomon, A.K., Dulvy, N.K., 2013. Ecosystem ecology: size-based constraints on the pyramids of life. Trends in Ecology & Evolution, 28(7), 423–431.

White, E.P., Enquist, B.J., Green, J.L., 2008. On estimating the exponent of power-law frequency distributions. Ecology, 89(4), 905–912.

Yurista, P.M., Yule, D.L., Balge, M., VanAlstine, J.D., Thompson, J.A., Gamble, A.E., Hrabik, T.R., Kelly, J.R., Stockwell, J.D., Vinson, M.R., 2014. A new look at the Lake Superior biomass size spectrum. Can. J. Fish. Aquat. Sci., 71(9), 1324–1333.

Yvon-Durocher, G., Montoya, J.M., Trimmer, M., Woodward, G., 2011. Warming alters the size spectrum and shifts the distribution of biomass in freshwater ecosystems. Glob. Change Biol., 17(4), 1681–1694.

## References

Andersen, K.H. and Beyer, J.E., 2006. Asymptotic size determines species abundance in the marine size spectrum. The American Naturalist, 168(1), pp.54–61.

Anderson, C.S., 1998. Partitioning total size selectivity of gill nets for walleye (Stizostedion vitreum) into encounter, contact, and retention components. Canadian Journal of Fisheries and Aquatic Sciences, 55(8), pp.1854–1863.

de Kerckhove DT, Milne S, Shuter BJ, Abrams PA. 2015. Ideal gas model adequately describes movement and school formation in a pelagic freshwater fish. Behavioral Ecology, 26(4): 1236–1247. https://doi.org/10.1093/beheco/arv073.

Edwards, A.M., Robinson, J.P., Blanchard, J.L., Baum, J.K. and Plank, M.J., 2020. Accounting for the bin structure of data removes bias when fitting size spectra. Marine Ecology Progress Series, 636, pp.19–33.

Edwards, A.M., Robinson, J.P., Plank, M.J., Baum, J.K. and Blanchard, J.L., 2017. Testing and recommending methods for fitting size spectra to data. Methods in Ecology and Evolution, 8(1), pp.57–67.

Gelman, A. and Rubin, D.B., 1992. Inference from iterative simulation using multiple sequences. Statistical science, 7(4), pp.457–472.

Kydd, J., H. Rajakaruna, E. Briski and S. Bailey. 2018. Examination of a high resolution laser optical plankton counter and FlowCAM for measuring plankton concentration and size. Journal of Sea Research, 133, pp. 2–10.

Lester, N.P., Sandstrom, S., de Kerckhove, D.T., Armstrong, K., Ball, H., Amos, J., Dunkley, T., Rawson, M., Addison, P., Dextrase, A. and Taillon, D., Wasylenko, B., Lennox III, P., Giacomini, H.C., Chu, C. 2021. Standardized Broad-Scale Management and Monitoring of Inland Lake Recreational Fisheries: An Overview of the Ontario Experience. Fisheries, 46(3), pp. 107–118.

Love, R.H. 1971. Dorsal aspect strength of an individual fish. J. Acoust. Soc. Am. 49, pp. 816–823

Mehner, T., Lischke, B., Scharnweber, K., Attermeyer, K., Brothers, S., Gaedke, U., Hilt, S. and Brucet, S., 2018. Empirical correspondence between trophic transfer efficiency in freshwater food webs and the slope of their size spectra. Ecology, 99(6), pp.1463–1472.

Millar R.B., 1992. Estimating the size-selectivity of fishing gear by conditioning on the total catch. J. Am. Stat. Assoc. 87, 962–968.

Millar, R.B., Holst, R., 1997. Estimation of gillnet and hook selectivity using log-linear models. ICES Journal of Marine Science 54 (3), 471–477.

Millar, R.B., Fryer, R.J., 1999. Estimating the size-selection curves of towed gears, traps, nets and hooks. Reviews in Fish Biology and Fisheries 9 (1), 89–116.

Neal, R.M., 2003. Slice sampling. Annals of statistics, pp.705–741.

Parker-Stetter SL, Rudstam LG, Sullivan PJ, Warner DM. 2009. Standard operating procedures for fisheries acoustic surveys in the Great Lakes. Great Lakes Fishery Commission Special Publication #09-01. Ann Arbor (MI).

R Core Team (2018). R: A language and environment for statistical computing. R Foundation for Statistical Computing, Vienna, Austria. URL https://www.R-project.org/.

Rudstam, L.G., Magnuson, J.J. and Tonn, W.M., 1984. Size selectivity of passive fishing gear: a correction for encounter probability applied to gill nets. Canadian journal of fisheries and aquatic sciences, 41(8), pp.1252–1255.

Sandstrom S, Rawson M, Lester N. 2013. Manual of Instructions for Broad-scale Fish Community Monitoring using North American (NA1) and Ontario Small Mesh (ON2) Gillnets. Ontario Ministry of Natural Resources. Peterborough, Ontario. Version 2013.2 35 p. + appendices.

Schneider, W.A., R.A. Reid, B.A. Locke and L.D. Scott. 1983. Studies of lakes and watersheds in Muskoka-Haliburton, Ontario: methodology (1976-82). Ont. Min. Envir. Data Report DR 83/1. 39 pp.

Smith, B.J., Blackwell, B.G., Wuellner, M.R., Graeb, B.D., Willis, D.W., 2017. Contact selectivity for four fish species sampled with north american standard gill nets. North American Journal of Fisheries Management 37 (1), 149–161.

Soule MA, Barange M, Hampton I. 1995. Evidence of bias in estimates of target strength obtained with a split-beam echo-sounder. ICES Journal of Marine Science, 52: 139–144.

Spiegelhalter, D., Best, N., Carlin, B. and van der Linde, A. (2002) Bayesian measures of model complexity and fit. Journal of the Royal Statistical Society, Series B, 64, 583–639.

Stauffer, H.B., 2007. Contemporary Bayesian and frequentist statistical research methods for natural resource scientists. John Wiley & Sons.

Tibbits, M.M., Groendyke, C., Haran, M. and Liechty, J.C., 2014. Automated factor slice sampling. Journal of Computational and Graphical Statistics, 23(2), pp.543–563.

Trebilco, R., Baum, J.K., Salomon, A.K. and Dulvy, N.K., 2013. Ecosystem ecology: size-based constraints on the pyramids of life. Trends in ecology & evolution, 28(7), pp.423–431.

Walker, S., Addison, P., Sandstrom, S.J., Lester, N.P., 2013. Contact retention selectivity of three types of gillnet gangs. Aquatic Research and Monitoring Section, Ministry of Natural Resources.

